# The Human-In-the-Loop Drug Design Framework with Equivariant Rectified Flow

**DOI:** 10.1101/2025.11.20.689585

**Authors:** Youming Zhao, Yanwen Huang, Xingang Peng, Haowei Lin, Qiao Xu, Yu-Zhe Shi, Xingchao Liu, Yijie Wang, Qiang Liu, Jian Peng, Jianzhu Ma

## Abstract

Advancements in AI-based drug design often face obstacles due to incomplete datasets, hindering progress to clinical trials. Human experts bring invaluable expertise and nuanced contextual understanding to drug design. There are two main difficulties in integrating human knowledge into the drug development process. First, human annotations are costly, and traditional machine learning algorithms require a large number of samples to be effective. Second, human experts are unable to accurately describe their expertise using natural language, nor can they precisely specify what kind of molecule is needed before seeing the generated molecules. To address these problems, we propose a new platform, called HIL-DD, for experts to infuse their experience by selecting molecules generated by AI that meet their criteria, or discarding those that do not. The core generative technology utilizes an Equivariant Rectified Flow Model (ERFM), which offers faster generation speeds than conventional diffusion models, enabling efficient human-AI collaboration. More importantly, we provide a user-friendly interface to ensure smooth and effective collaboration between human experts and AI systems. Rigorous experiments demonstrate that our system can produce 3D molecules that align with expert expectations in minimal interactive sessions. These generated molecules maintain drug-like qualities comparable to those created by current state-of-the-art models.

---

Despite significant advancements in using AI to aid drug design, only a few AI-designed drugs have progressed to clinical trials, and none have received approval from the U.S. Food and Drug Administration (FDA) thus far. The reasons behind this dilemma are complex. First, AI algorithms heavily rely on data, and the lack of comprehensive data hampers their ability to generate accurate predictions. Second, the biological systems involved in drug development are multifaceted, making it difficult for AI models to capture all relevant complexities with a single model. Traditionally, human experts play a crucial role in designing and optimizing drug candidates based on their expertise and understanding of biological systems. However, human-driven processes can be time-consuming, expensive, and limited by the availability of expertise. Therefore, a natural question arises: can we integrate human wisdom and experience with existing AI systems to design drug candidates more efficiently? To address this problem, we developed a new Human-In-the-Loop Drug Design (HIL-DD) system, facilitating direct interaction between human experts and AI for codesigning 3D molecules. This work focuses on generating 3D molecules to bind a protein pocket based on its 3D structure, but the approach can be generalized to other AI drug design tasks. Initially, AI generates candidate 3D molecules that can fit the protein pocket based on predefined objective functions. These candidates are then presented to human experts for evaluation and selection based on their perceived promise and favorability. HIL-DD utilizes this feedback to fine-tune its parameters, improving its generation process in subsequent rounds. By enabling collaboration between humans and machines, HIL-DD combines the analytical power of AI with the creativity, experience, and deep contextual understanding of human pharmacists.

The most common way of human-machine interaction (HMI) is for human annotators to rank or score the outputs generated by machine learning models [38, 26, 17, 1, 10, 20].

The implementation of the HMI system presents new challenges to the generative models. First, we expect the generative model to respond promptly to feedback from human experts. Conventional diffusion models, although powerful, often require excessive inference time to generate a high-quality sample. Second, the generative model needs to effectively capture the preferences of human experts during the molecule selection process with only a few human annotations. To meet these requirements, we employed a recent advancement in generative models, named Rectified Flow (RF), as the backbone model of the HIL-DD system [18, 19]. The RF model can generate samples in a few steps, such as 10, because it learns an ordinary differential equation (ODE) that follows a straight path. The main advantage of the RF model over diffusion models is its faster generation speed while maintaining expressive power, making it more suitable for interacting with human experts. We further extended the RF model to an Equivariant Rectified Flow Model (ERFM) to capture the rotational and translational invariance properties of 3D molecules. To enable a fast response to human feedback, we also introduced multiple new loss functions that drive the straight line toward the generation of positive molecules selected by human experts compared to the negative ones.

Experimental results show that our method can effectively infer human intentions from limited interactions with the preference oracle and demonstrate robust error tolerance against noisy feedback. Another finding is that our system, while optimizing for human preferences, does not compromise on other pharmacological indicators, remaining on par with other state-of-the-art 3D molecule generation approaches. We observed that not all metrics could be continuously improved with an increased number of expert annotations. The ultimate performance of the human-in-the-loop design process depends on: 1) the inherent complexity of different metrics themselves, and 2) whether the model can sample molecules that meet human expert expectations.

## Results

### Overview of the HIL-DD framework

Due to the high cost of human annotations, we employed a simulator to evaluate the performance of HIL-DD, which plays the role of human experts, to provide preference annotations to our system. In each experiment, the simulator focuses on a specific metric which is strongly associated with the potential of a molecule to possess drug-like properties. We trained a generative model based on the ERFM on the CrossDocked training data [8]. In each round of interaction, the model generates *N* molecules, then the expert simulator selects *M* (0 ≤ *M* ≤ *N*) molecules from them based on its own preferences (Figure 1b). It is possible for the simulator to either select all molecules or not select any molecule at all. Next, the model takes new preference data as inputs and updates its parameters with additional loss functions. Note that the ERFM is not aware of the underlying metric on which we focus throughout the fine-tuning process. Last, we evaluated the improvement of metric compared to the original model with respect to different numbers of preference annotations.

**Fig. 1.**
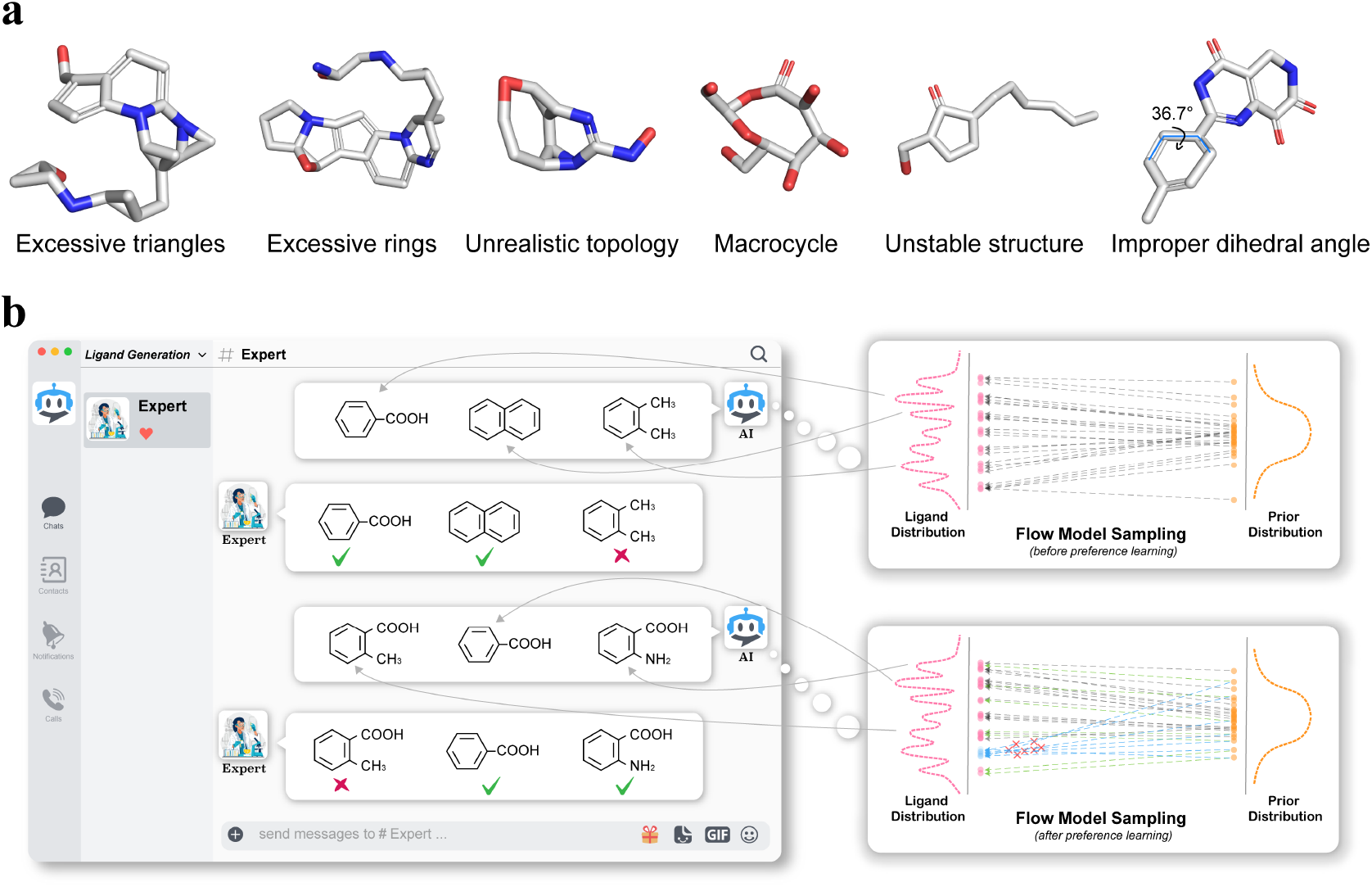
Molecules with obvious structural defects and HIL-DD system structure. **(a)** Molecules generated by popular 3D molecule generation methods, showing obvious structural defects. **(b)** Overall framework of HIL-DD. AI generates multiple 3D molecules, and human experts select some of these molecules based on their preferences. AI then learns from human preferences and continues to generate the next round of 3D molecules.

Overall, we have designed two types of loss functions to make the molecules generated in the next round more aligned with the expert preferences. One type is the **preference loss function** which attempts to directly modify the direction of the straight lines generated by the ERFM model, so that they are more inclined towards the samples selected by the experts. The second type is the **guidance loss function** which is equivalent to training a binary classifier using the expert-provided preferences to predict whether a sample represents an expert preference. The gradients of this binary classifier with respect to the samples can then provide guidance for the generation of molecules in the next round. For the preference loss function, we further developed multiple alternative forms of loss functions including cross entropy loss, binary cross entropy loss, and hinge loss (Figure 2). Note that these loss functions are not mutually exclusive and can be used together. More technical details could be found in the **Methods** section.

**Fig. 2.**
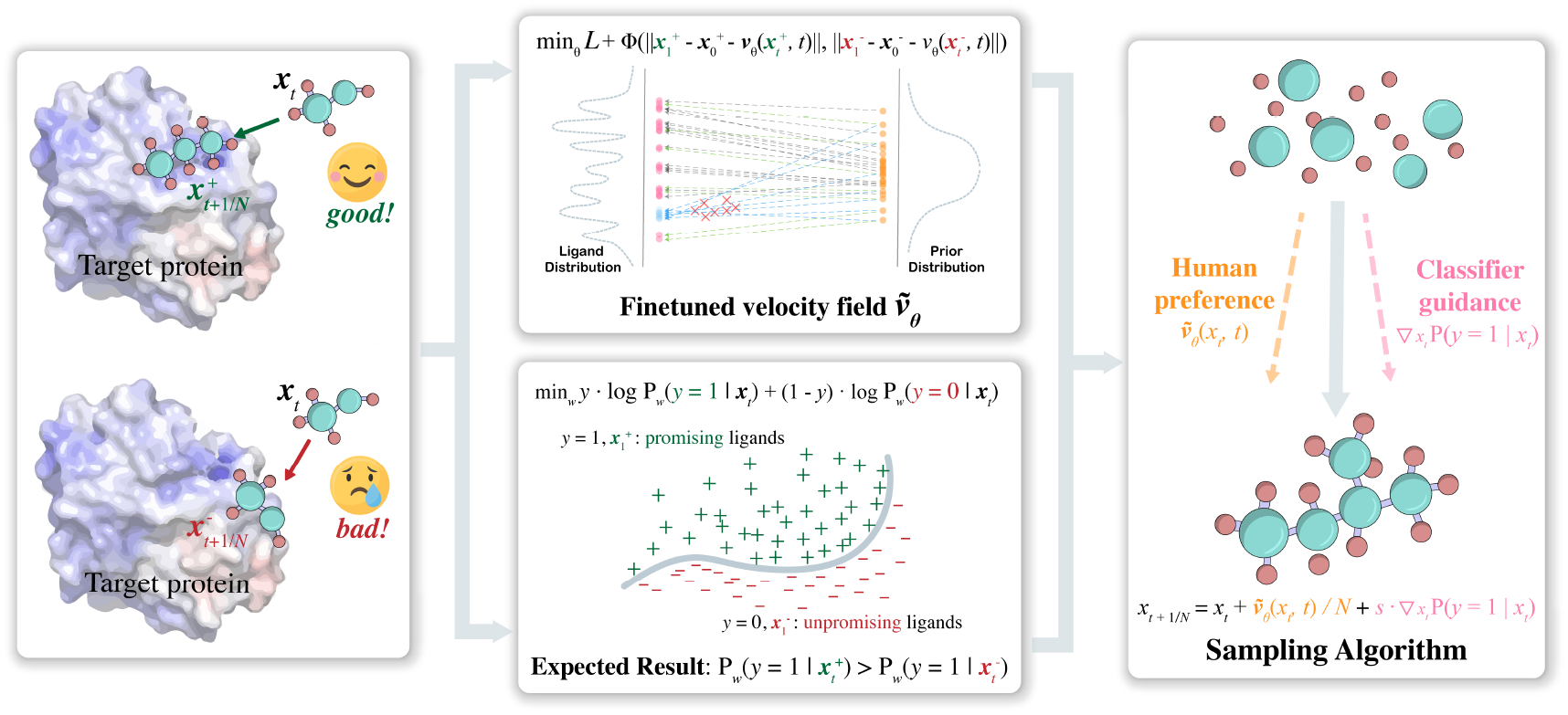
The schematic diagram of the mathematical principles of preference learning in HIL-DD.

### Performance of 3D molecule generation

We first evaluated the quality of 3D molecules generated by ERFM based on protein 3D pockets before and after using human feedback (Figure 3). ERFM represents the performance without using any expert annotated data, and the HIL-DD represents the performance after learning from human expert annotations. The details about running HIL-DD are depicted in Supplementary, section 5-7. The performance was evaluated on 100 protein pockets of the test set. We compared our method with the state-of-the-art 3D molecule generation models AR [23], Pocket2Mol [29], and TargetDiff [11]. The technical details are presented in the **Methods** section. The hyper-parameter selection and model architectures could be found in the Supplementary, section 5-6. The implementations of baseline models are described in the **Methods** section.

**Fig. 3.**
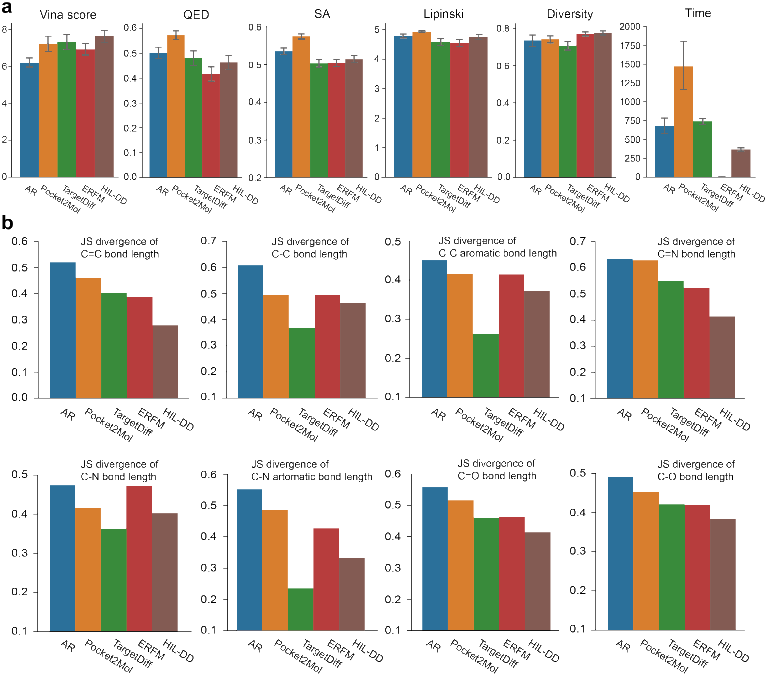
Performance of 3D molecule generation. **a**. Summary of molecular properties related to the drug-likeness of the 3D molecules. The mean values and 95% confidence intervals are reported for 100 protein pockets from the test set with 100 samples for each pocket. **b**. Jensen-Shannon (JS) divergence between the distributions of bond distance of training data and the generated molecules. A lower value is better. Here, the bars for ERFM are produced via molecules generated by ERFM with 10-step sampling.

We have carefully chosen a set of widely adopted metrics to thoroughly evaluate the characteristics of the sampled molecules. These metrics serve as reliable indicators for assessing the quality and potential of the generated molecules including: 1) Vina score: an estimation of the binding affinity between the molecules and the protein, indicating the potential therapeutic effect as drug candidates; 2) QED (Quantitative Estimation of Drug-likeness), which quantitatively evaluates the drug-likeness of a molecule; 3) SA (Synthetic Accessibility) score: a measure of the synthetic accessibility of a molecule, ranging from 0 to 1, with higher value indicating greater ease of synthesis; 4) Lipinski: Lipinski’s Rule of Five is an empirical guideline used to assess the drug-likeness of molecules. Compliance with this is often considered favorable for drug candidates; 5) Diversity: a measure of the variation within the generated set of molecules. It is calculated as the average Tanimoto similarity among the molecules for individual protein pockets; 6) Time: the time duration required to generate 100 molecules for a specific protein pocket. It provides insights into the efficiency and speed of the generation process of our ERFM. By considering these comprehensive metrics, we can holistically evaluate the potential drug candidates and make informed decisions regarding their viability for further development and testing.

As shown in Figure 3, our model has demonstrated comparable performance compared to all the existing methods across all the evaluated metrics. This is also the first work which adopts the Rectified Flow model on the task of 3D molecule generation conditioned on protein pockets. It is worth noting that ERFM can generate 3D molecules with high binding affinity for protein pockets, and these protein pockets have never appeared in the training set. When considering the generation time, our ERFM with 10 steps significantly outperforms other diffusion-based methods, such as TargetDiff which need 1,000 steps to generate high quality molecules. This efficiency is critical in the scenario of human-in-the-loop drug design, where timely decision-making and iterative processes are important. This shows that ERFM can act as a highly promising generative model, offering great potential for accelerating drug discovery.

### Performance of the HIL-DD framework with simulated preferences

Next, we evaluated the performance of the model under different human preference choices. We randomly selected five proteins (PDBID: 2V3R, 14GS, 4YHJ, 5W2G, 1COY) as the test proteins and sampled 26,000 molecules for each protein. We chose promising and unpromising molecules from the 26,000 samples. The details about selecting samples can be found in the Supplementary, section 7. Empirically, we found that models with different preference loss functions perform similarly across different protein pockets under the same human preferences. However, when given the same protein pocket, the model’s performance varies depending on different human preferences. Therefore, we focused on comparing the performance of different human preferences for the same protein. Figure 4 displays the experimental results of protein 2V3R under different types and numbers of preferences, the results for other protein pockets are presented in the Supplementary, section 3.

**Fig. 4.**
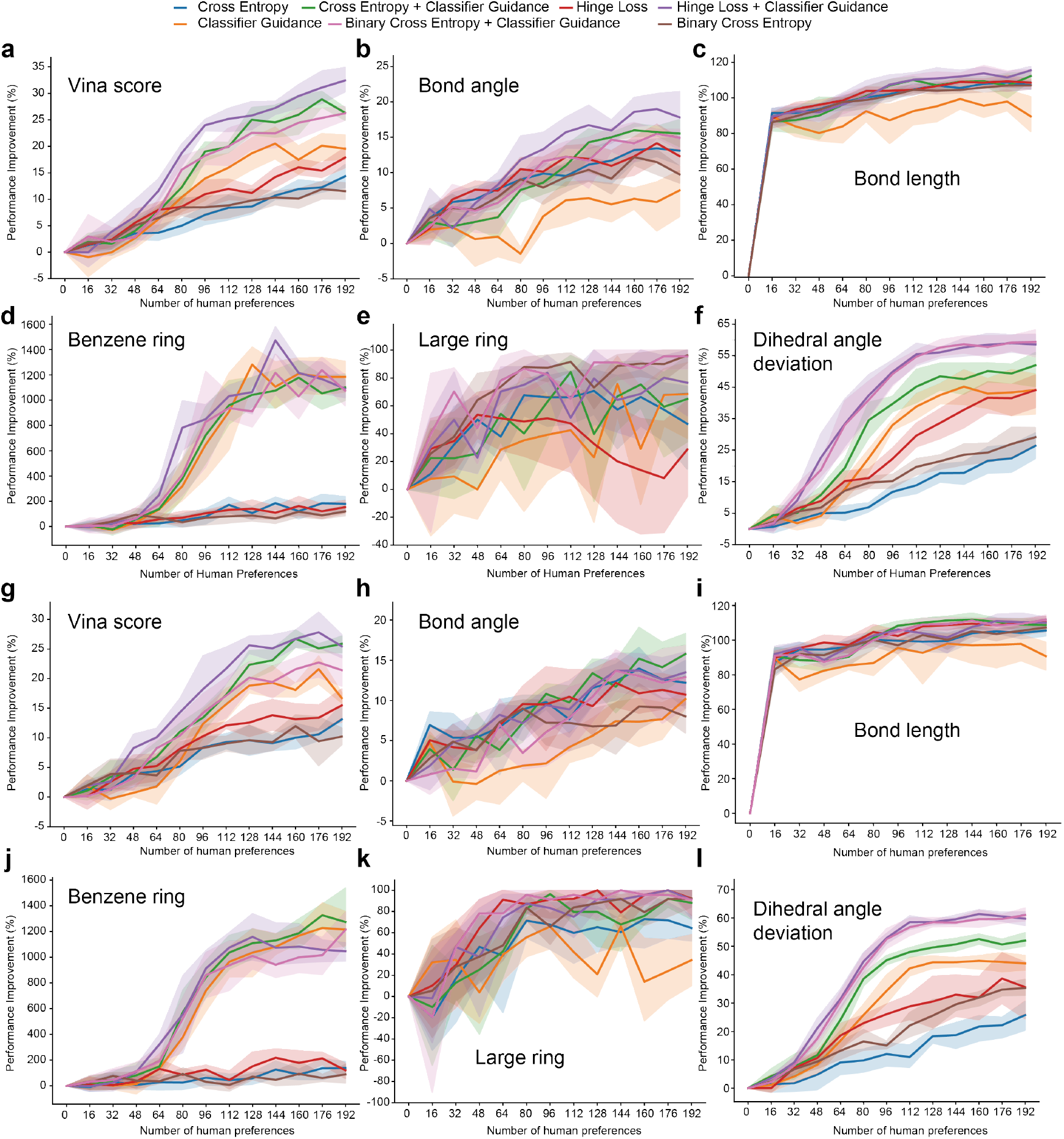
Performance of the HIL-DD framework. We measure whether the generated molecules reflect the preferences of human experts. For instance, if human experts select molecules with high Vina scores, we assess whether the Vina scores of newly generated molecules are improved. **a** to **f** represents the performance of the incremental annotation of samples by human experts. AI learns in an online learning fashion. **g** to **l** represents AI learns all the annotated data at once.

The first human preference we considered was the binding affinity between a protein pocket and the generated 3D molecules. Even human experts might not have an instinct about the binding affinity but they can invoke a third-party tool to calculate it. We adopted the Vina docking score implemented by Quick Vina 2 [12] to approximate the binding affinity, while others could certainly use other tools, such as AutoDock Vina [39], GOLD [40], Glide [9], Schrödinger Suite [34], MOE [41], DOCK [4], etc. Vina docking score can only act as a preference because it is hard to be directly incorporated into the generative model because it optimizes a knowledge-based energy function based on multiple physical and statistical terms among atoms and chemical bonds. The average Vina docking score achieved by ERFM without any feedback is −7.46, which is calculated over 26,000 sampled molecules for the protein pocket of interest. As shown in Figure 4a, preference loss functions (the brown, red and blue lines) can utilize the expert feedback and all converge at the level of 10% of performance gain. Classifier guidance loss function (the orange line) is effective when very few preferences are provided but stop very early at the level of 18% of performance improvement. Further improvement to 32.43% (the purple line), which corresponds to a Vina docking score of −9.88, is observed when two types of loss functions are combined together (the purple, green and pink lines). For this protein, the Vina score for the native complex structure is −10, so our model is already approaching the upper bound of data with only 96 positive and 96 negative samples that are annotated.

The second type of preference encompasses bond angle and bond length. The reason we choose chemical bonds as the preference is that most existing 3D molecular generation models, especially diffusion models, overlook chemical bonds during training and can only add them back using post-processing software after generating atoms [13, 11]. This methodology often leads to discrepancies in both bond lengths and bond angles compared to their distributions in the training set [28]. The first and third molecules in Figure 1a are typical instances of errors caused by incorrect addition of chemical bonds in this post-processing way. In this experiment, we evaluated the quality of the generated molecules based on the log-likelihood of the chemical bond distribution. Specifically, we assessed whether the bond lengths and bond angles of the generated molecules match the distributions observed in the training data using the log-likelihood. As shown in Figures 4b and 4c, it is easy to improve the performance of bond length generation in comparison to the bond angle using preferences. HIL-DD could achieve ∼ 100% of log-likelihood improvement with preference learning. Bond angle is more challenging to learn because patterns of bond angles are more complex than bond lengths as they involve more atoms, and thus requires a larger amount of data for accurate learning. However, the combination of the hinge loss function and the classifier guidance still achieves a maximum improvement of 18.99% when preferring samples with desired bond angles.

Next, we have found that current molecule generation models tend to produce incorrect distributions of cyclic compounds of different sizes. For instance, Guan et al. [11] shows their method produces a large proportion of 7-membered and 8-memebered rings which have been proved to be rare in the training data. This observation is easy to be detected but hard to be captured as constraints in the generative model. In the next experiment, we studied whether HIL-DD could correct such bias based on limited human annotations and preference learning. Empirically, the simulator prefers the generated molecules with 3D structures of benzene rings that have a quality above a certain threshold (**Methods**). As shown in Figure 4d, before applying preference learning, 2.78% of the generated molecules contain benzene rings, which is significantly lower than 59.86% in the training set. With 192 preference selections, this ratio is improved by 1471.67%, which indicates that HIL-DD can generate 43.69% molecules containing qualified benzene rings.

Another drawback of the current generative model is that they tend to produce cyclic compounds that are either too large or too small (Figure 1a). We investigated whether experts could inform the generative model about the tendency to avoid generating molecules with excessively large cyclic structures (≥ 9 atoms in our implementations) through limited annotations. As shown in Figure 4e, the results are similar to the previous benzene ring experiment where the HIL-DD system can effectively utilize human annotations to reduce the presence of large cyclic structures in the generated molecules.

The last metric of interest is dihedral angle deviation. Dihedral angle measures the angle between two planes formed by three bonds, which is an important metric to assess whether generated molecules satisfy certain geometric constraints. For instance, the six atoms of a benzene ring should lie on a common plane. The average dihedral angle of the ERFM model is 17.13^*◦*^. To improve this metric, the simulator provides

AI with samples that have very low average dihedral angle deviations (≤ 5^*◦*^ in this case). After applying preference learning, the fine-tuned model improves this metric by 59.34% as shown in Figure 4f, which indicates the model can generate more realistic molecules in terms of chemical and geometric constraints.

Expert annotations were added sequentially in the above experiments, allowing experts to stop generating new molecules once they were satisfied with the results. Another interesting strategy to explore is having experts annotate the same number of samples at once, allowing the model to learn from these annotated molecules collectively. As shown in Figures 4g-l, the performance impact on the model from learning the same number of samples in a single batch compared to sequential learning is marginal. Empirically, we did not adopt this strategy as it is relatively time-consuming and cognitively demanding for human experts. Some typical molecules generated by ERFM and HIL-DD w.r.t. the preferences of Vina score, bond angle, and benzene ring are demonstrated in Figure 5. The molecule visualization for the remaining preferences can be found in Supplementary, section 10.

**Fig. 5.**
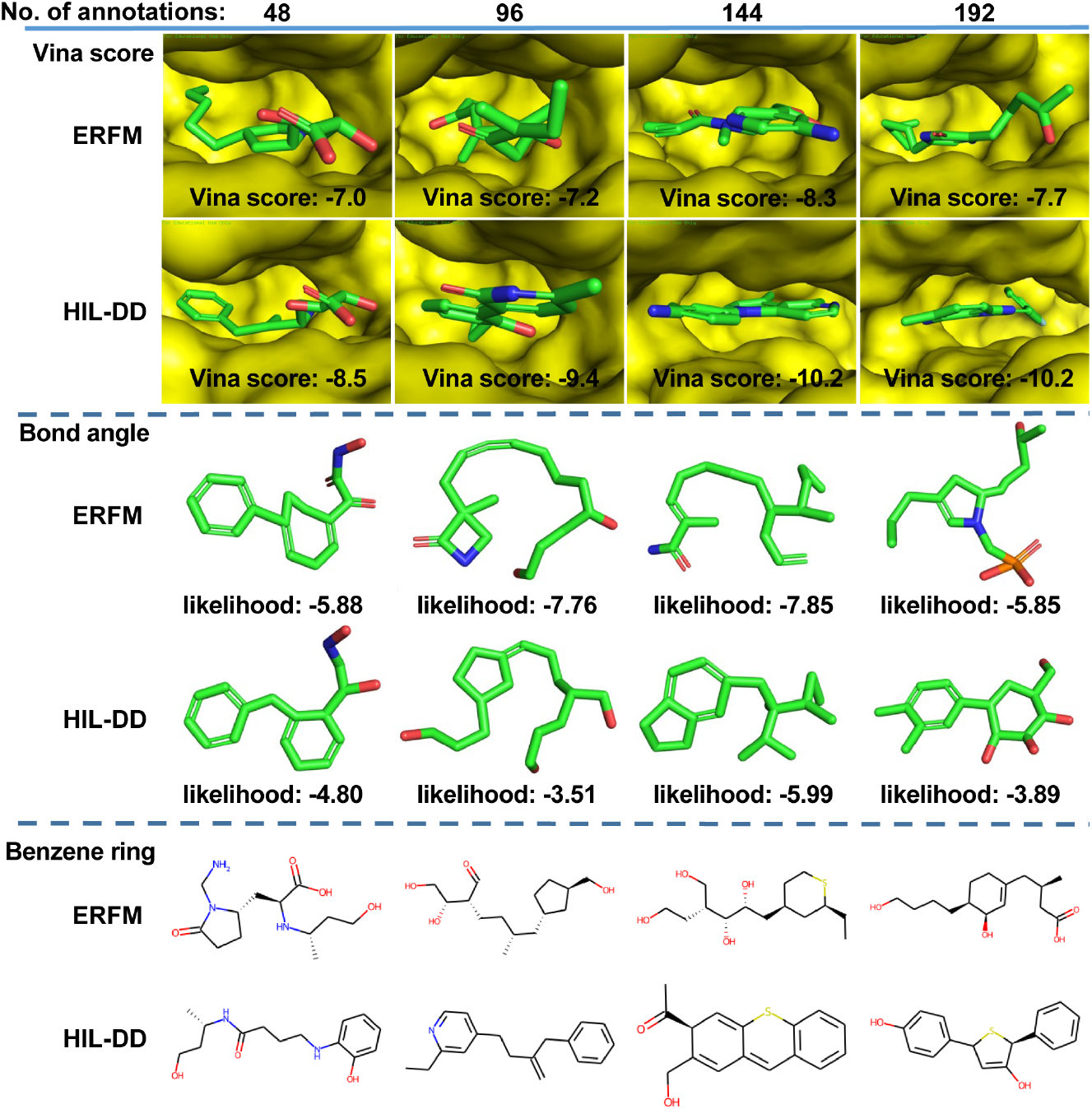
Drug candidates generated by ERFM and HIL-DD. The molecules presented in four columns are generated by fine-tuning the ERFM with 48, 96, 144, and 192 human preferences, respectively. The likelihood is measured in a log scale.

### Performance of the HIL-DD with human preferences

To further evaluate our algorithm, we invited five medicinal chemists from both academia and industry to provide preferences to the prediction of our model. The drug target we select is Epidermal Growth Factor Receptor (EGFR), which is involved in multiple biological processes such as cell proliferation, survival, adhesion, migration and differentiation [37, 31]. Mutations of EGFR contributes to the uncontrolled growth and division of cells, and ultimately drive transformation to non-small cell lung cancer. We selected this EGFR to study our human-machine-interaction system for two reasons. First, the FDA has granted approval to multiple small-molecule drugs targeting this gene, hence medicinal chemists have a certain level of understanding about which types of molecules might inhibit EGFR [15, 22]. Second, most patients develop drug resistance to nearly all the EGFR inhibitors after 1–2 years of treatment, which poses a major challenge in clinical practice. As half of these drug resistance cases are caused by the secondary EGFR T790M mutation, we chose the crystal structure of EGFR with the T790M mutation (PDB ID: 6JX0) to generate 3D molecules. As the FDA-approved drugs all contain the phenylaminopyrimidine substructure, at the first round human experts selected 199 molecules with the phenylaminopyrimidine substructure. The experts then examined these molecules and selected eight molecules as drug candidates based on their experience. Due to the presence of multiple experts, each expert eliminated the molecules that they considered to have an unreasonable structure or were difficult to synthesize. The HIL-DD model learned the human preferences and generated another 75 molecules. With the same criteria, the experts identified five drug candidates which met all their criteria.

We carefully examined these two rounds of molecules. As shown in Figure 6a, the preference loss decreased from 3.5 to 3.36, indicating the model learned certain knowledge from the human preference, which drove the success rate defined by the experts to increase from 8/199 to 5/75 by 65.83% (Figure 6b). Next, we interviewed and summarized the main criteria adopted by the experts and examined whether these criteria were more satisfied in the second round. Based on the experts’ feedback, the initial model had the tendency to generate chemically infeasible macrocyclic molecules and fused ring structures. Although well-designed macrocycles might offer additional binding affinity and stability, the macrocycles generated by the model has been identified to have inappropriate sizes and shapes, and thus are less likely to bind the target. Another characteristic of these macrocyclic molecules that concerns the experts is the complex linker structure, relating to poor biological activities and substantial synthetic costs. The initial model exhibited a tendency to generate an unexpected number of molecules featuring uncommon fused ring structures that had not been previously reported. As shown in Figure 6c top, the irregular double bond in the fused ring could render the molecule unstable and highly reactive at room temperature, potentially leading to toxicity. The size and rigidity of such fused ring structures raise concerns about solubility and drug-likeness. Figures 6d and 6e demonstrates that the model generated less molecules with the unfavorable macrocyclic and fused ring structures identified and excluded in the selection phase, validating the effectiveness of the inclusion of expert preference. On the other hand, experts showed strong interests in the generated molecules that exhibited significant modifications to the pyrimidine moiety, especially ring-jointing modifications (Figure 6c bottom). This preference was motivated and supported by a series of published work [43, 36, 27], suggesting the potential benefits in targeting the EGFR T790M mutation. As shown in Figure 6f, the resultant model exhibited an increase in the occurrence of these structurally modified compounds. In the end, we observed that several molecules generated after the preference learning contained similar substructures as currently in clinical or FDA-approved drugs targeting EGFR T790M mutations, suggesting that these compounds may be effective against the EGFR T790M mutations.

**Fig. 6.**
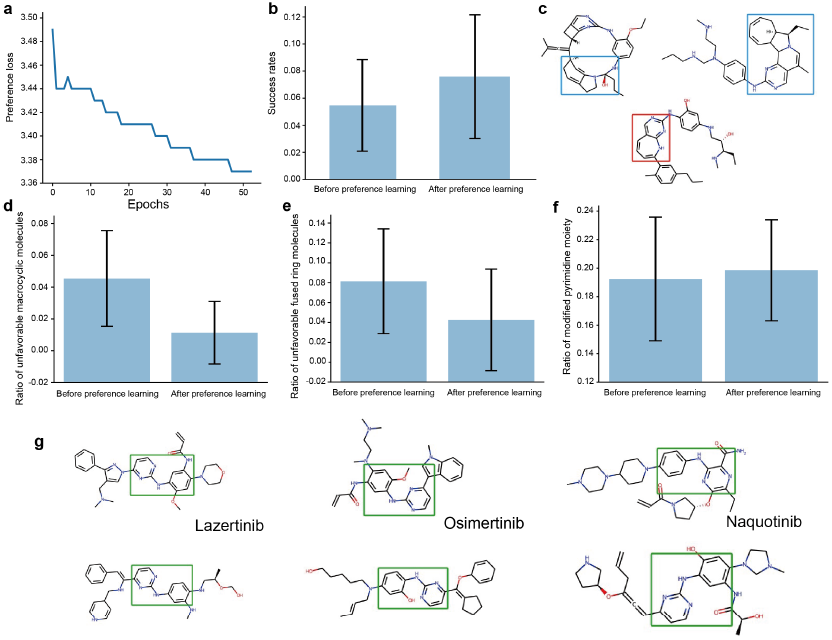
Performance of the HIL-DD with human preference. **a**. The loss function of preference learning learned from human annotations. **b**. Success rates of generating molecules satisfying the human experts before and after applying the preference learning. **c**. Typical molecules with unfavorable (blue) and favorable (red) substructures. **d-f**. Ratio of unfavorable macrocyclic molecules, unfavorable fused ring molecules, and modified pyrimidine moiety before and after the preference learning. **g**. HIL-DD model identified molecules similar to FDA-approved drugs. The error bars represent standard deviations on 5 independent runs.

### Graphical user interface of HIL-DD

As shown in Figure 7, to efficiently communicate with human experts, we developed a lightweight and decentralized Graphical User Interface (GUI) based on Vue3 (https://vuejs.org/) setup and webpack bundling (https://webpack.js.org/). The software is optimally designed for local hosting, thereby facilitating integration with existing computational resources. The software leverages a real-time push mechanism, dynamically providing a list of molecules for human experts to select. Human experts can provide their preference through three distinct options: “Like”, “Dislike”, and “Preview”. Upon selection of a molecule, its metrics are shown instantaneously updated and its structure is rendered for immediate feedback, which exemplifies the interactive design of our software.

**Fig. 7.**
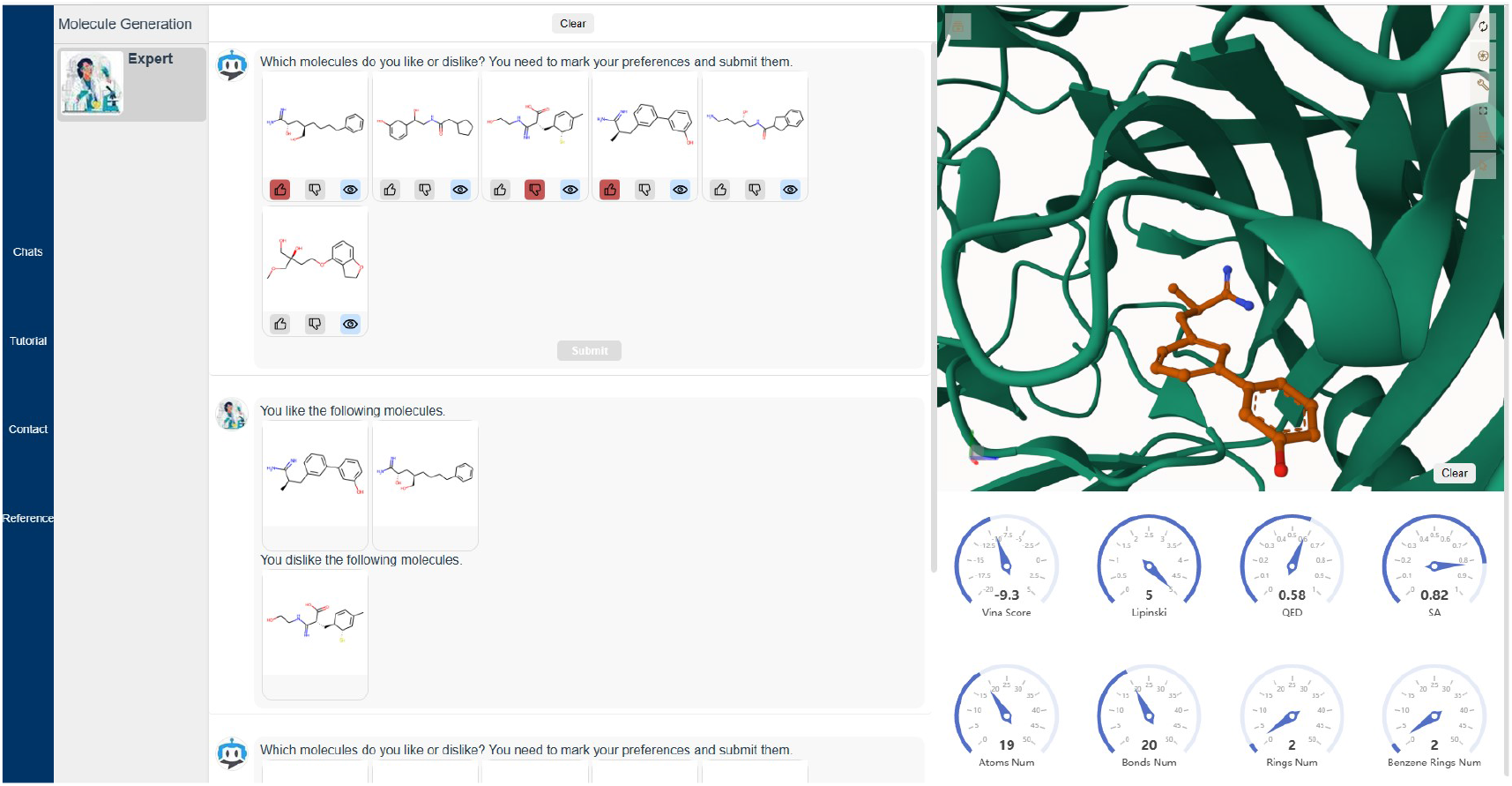
Graphical user interface of HIL-DD

The front end features: 3D molecular structure rendering is implemented by the open source project Mol* [35], which allows multiple fundamental operations on the 3D molecules. 2D molecular structures are rendered using rdkit.js, providing users with a comprehensive understanding of molecular structures. D3.js is employed to generate interactive diagrams [2].

## DISCUSSION

The HIL-DD framework can be classified within the realm of active learning. However, in comparison to conventional active learning methodologies, HIL-DD presents multiple distinctive characteristics. First, current active learning approaches are commonly employed in a supervised learning setting to predict sample labels. They primarily emphasize assessing the confidence of label predictions, whereby human experts need to annotate samples with low confidence to facilitate subsequent model retraining. Conversely, the HIL-DD incorporates an unsupervised generative model as the machine learning module and does not need to calculate the confidence of the predictions. The inherent challenge of generative models lies in the absence of any predefined labels. Consequently, within the traditional active learning framework, human experts will encounter difficulties in determining how to annotate such samples. To address this problem, we adopt a preference learning methodology, wherein human experts can provide suggestions to AI models by using sample comparisons. Second, with regard to the learning process, traditional active learning necessitates model retraining following the acquisition of new labeled data with the original model fixed. In our framework, we fine-tuned the trained generative model by introducing multiple novel preference loss functions.

Another machine learning model related to ours is a recently emerged chatbots based on Large Language Models (LLM) which are also generative models and need human feedback. The LLM-based chatbots, such as GPT [30, 3], PaLM [6] and LLAMA [24], all can quickly adjust their behavior in response to human feedback. In drug design, there have been works developed to combine LLMs with molecular generation, attempting to allow experts to manipulate molecule generation using natural language [25, 5, 14]. Experts can modify specific chemical groups through natural language instructions. However, the reason medicinal chemists require computer assistance is not because they cannot accurately draw the relative positions of benzene rings inside the protein pockets. It is because they cannot quantitatively describe their experience in drug development in a mathematical form. The core contribution of our HIL-DD system is to provide a novel way of interaction between experts and AI models, to extract human knowledge and expertise to correct the bias introduced by the computational model.

Our methodology is not only limited to rectified flow models, but it can also be readily applied to other ODE-based generative models, as well as to a broader range of deterministically generative models. Specifically, by utilizing a pretrained generative model, we have the flexibility to adjust the regular loss for positive and negative samples during the fine-tuning process. This is achievable by incorporating the preference loss functions introduced in **Methods**. This approach allows us to reduce the regular loss for positive samples and increase it for negative samples. Therefore, our methodology can be applied effectively to various generative models by adapting and incorporating these preference loss functions into the fine-tuning stage.

Our work focuses on the optimization of molecules based on a single objective, while human experts often evaluate molecules by considering multiple criteria in a holistic manner. Although human experts can utilize the current HIL-DD system to preferentially select molecules based on one criterion and then proceed to a second criterion, this approach undoubtedly reduces efficiency. The reason is that the samples that emerge from the selection based on the first criterion may exhibit favorable or unfavorable features for the second criterion. Therefore, in the future work, we will focus on the multi-objective optimization of 3D molecules.

## Methods

### Equivariant neural network

Since we need to model the 3D structures of both small molecules and proteins, it is essential to consider the rotational and translational invariance of the neural network model. A function *f*: ℝ^3^ → ℝ^3^ is E(3) equivariant if *f* (***Rx*** + ***t***) = ***R****f* (***x***) + ***t*** for any orthogonal matrix **R** ∈ ℝ^3*×*3^ and any column vectors ***x, t*** ∈ ℝ^3^. *f* is considered as *O*(3) equivariant if *f* (***Rx***) = ***R****f* (***x***) and translation invariant if *f* (***x***+***t***) = *f* (***x***)+***t***. A function *f* is *E*(*n*) equivariant if it is equivariant with respect to rotations, translations, reflections and permutations in an *n*-dimensional space. In this work, we employed *E*(*n*) Equivariant Graph Neural Networks (EGNNs) to model a protein and a small molecule pair as one graph [32], in which atoms are nodes and each atom is connected to its *K* nearest atoms in the Euclidean space (*K*=48 in our implementations). The layers of the EGNN model are defined as follows,

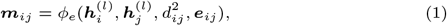

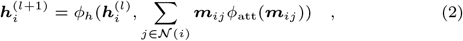

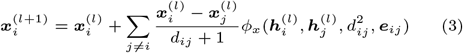

Here, ***x***_*i*_ and ***h***_*i*_ denote the 3D coordinate and hidden representation for atom *i*, respectively. ***m***_*ij*_ and ***e***_*ij*_ denote the hidden representation and feature vector for edge *i, j*, respectively. *d*_*ij*_ is the distance between ***x***_*i*_ and ***x***_*j*_. *ϕ*_*e*_, *ϕ*_*h*_, *ϕ*_att_, and *ϕ*_*x*_ are all neural network models. Note that the Center of Mass (COM) of the protein atoms is shifted to 0 to preserve translational invariance. This technique is commonly referred to as the subspace trick [16, 42, 13, 33]. For a detailed explanation, please refer to the Supplementary, section 1.

To better capture the bond information, we added the bond representation ***b***_*ij*_ and its update rules in the EGNN model as follows,

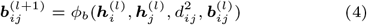

where *ϕ*_*b*_ is also a neural network model, and atom *i* and atom *j* form a chemical bond. Eq. (4) indicates that learning bond representation can influence the learning of atom representations. Based on the ablation study in the Supplementary, section 4.7, the quality of the generated 3D molecules with bond neural network is higher than the model trained without bond neural network. The network architecture of *ϕ*_*e*_, *ϕ*_*b*_, *ϕ*_att_, *ϕ*_*x*_, and *ϕ*_*b*_ could be found in the Supplementary, section 4.

### Equivariant rectified flow model (ERFM)

Following previous works [19, 18], we constructed an ODE process, which transports a tractable prior distribution to the data distribution conditioned on a particular protein pocket. We modeled this transport process as a Rectified Flow (RF) model and used a neural network to fit the flow as follows,

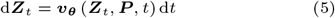

where ***v***_***θ***_ is the flow neural network using EGNN as the backbone, and ***Z***_*t*_ = (1 −*t*)***Z***_0_ + *t****Z***_1_ with *t* ∈ [0, 1]. ***Z***_*t*_ denotes the generated 3D molecule at time stamp *t*. ***P*** denotes the representation of the protein pocket which contains multiple atom coordinates and other features. Note that ***P*** is fixed during the whole transporting procedure, so we omit ***P*** to keep the notation uncluttered hereafter. The features of protein pockets could be found in the Supplementary, section 4.

As discussed in the previous section, our goal is to learn a function ***v***_***θ***_ which maps a sample ***X***_0_ from Gaussian distribution to another sample ***X***_1_ from the observed data distribution. Intuitively, given an intermediate state ***X***_*t*_, the optimal direction that transports ***X***_*t*_ towards ***X***_1_ is ***X***_1_ − ***X***_0_. Thus, the key concept of our optimization process is to optimize the velocity field ***v***_***θ***_(***X***_*t*_, *t*) and make it close to the optimal direction ***X***_1_ − ***X***_0_. The training objective function could be written as,

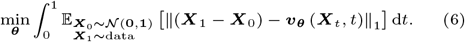

To solve (6), in each epoch we sampled a random pair ***X*** = (***X***_0_, ***X***_1_) and then optimized the following equivalent objective function,

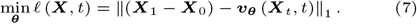

where *t* ∈ [0, 1] and ***X***_*t*_ = *t****X***_1_ + (1 − *t*)***X***_0_. Note that *ℓ* (***X***, *t*) is also called the straightness at *t* [19]. We optimized (7) using AdamW [21], a popular stochastic gradient descent algorithm. Hyper-parameters related to the optimizer can be found in the Supplementary, section 5. Gradients with respect to model parameters are computed by standard back-propagation. Once the function ***v***_***θ***_ is trained, 3D molecules can be generated by iteratively running the following Euler steps given a pocket ***P*** as follows,

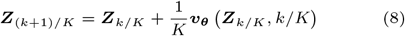

Here *K* is the number of time steps for training and *k* = 0, 1, · · ·, *K* − 1. After running *K* steps, we get a 3D small molecule ***Z***_1_ which is supposed to bind to the given pocket ***P*** with high binding affinity. Since the pocket of interest is taken from the CrossDocked dataset, the number of atoms of any generated molecule is the same as the reference molecule for the pocket. Detailed sampling procedure is described in the Supplementary, section 9. Note that the conventional diffusion model can also be formulated in the form of (5). The key difference between the ERFM model and the conventional diffusion model lies in the generation process. For the ERFM model, all randomness occurs in the initial step, where the function ***v***_***θ***_ transforms a random sample into a molecule in 10 steps, due to the easy optimization objective of learning straight paths. The traditional diffusion model requires much more steps, typically 1000, with each step making minor changes to the sample and incorporating a certain amount of randomness, which generates samples in a relatively cautious and incremental manner. However, in the context of human interaction, we need to rapidly generate a large number of samples and adapt the model behavior quickly based on human preferences. Therefore, the generation approach of the ERFM is more suitable for this purpose.

### Preference learning

In the HIL-DD framework, human experts communicate with the AI by providing their preferences for the molecules generated by AI. These preferences could be provided due to HIL-DD generating a molecule that meets the criterion set by the experts, or some of the rejected molecules exhibit obvious (8) errors. Therefore, the most crucial module in HIL-DD is to model user preferences and to train the model which can rapidly adjust the molecules it generates based on a limited number of preferences. The intuition of our solution is to “promote good and restrain evil”, i.e. to enhance the probability of generating the selected molecules and decrease the probability of generating the deselected molecules. Technically, the neural network function ***v*** should predict straight lines that map our randomly generated sample to the samples selected by human experts instead of mapping to the samples not chosen by human experts. More formally, let *π*_0_ denote the Gaussian distribution and *π*_1_ denote the data distribution. We used 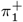 and 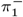 to represent distributions for the selected and unselected molecules based on the preference of human experts, respectively. For any interpolant 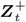 between 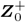 drawn from *π*_0_ and 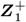 drawn from 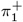, minimizing the designed objective function pulls the straight line 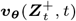 to be close to the vector 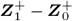, and at the same time pushes 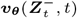 to be far away from 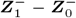. In this case, 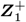 acts as a source of attraction, pulling the line towards it, while 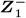 acts as a source of repulsion, pushing the line away from it. Including human preference 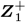 and 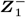 is equivalent to shrinking the space of 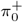 and 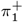 while expanding the space of 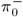 and 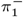. This is shown empirically by a toy experiment in the Supplementary, section 2.

#### Loss function

If a human expert selects *N* molecules based on their preferences from a total of *M* molecules, we then collect a total of *N* (*M* − *N*) pairs of positive and negative samples. Given a pair of these samples conditioned on a pocket ***P***, we can optimize the following objective function,

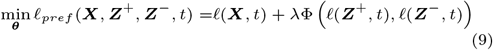

where *t* ∈ [0, 1], *λ* is a hyper-parameter, ***Z*** = (***Z***_0_, ***Z***_1_), and *ℓ*(·) is defined in (7). Φ(*a, b*) is a function designed to decrease the value of *a* and increase the value of *b* during the optimization process. We have provided three alternative functions of Φ and carefully studied their performance in the **Results** section.

#### Cross entropy loss

The cross entropy function is defined as follows,

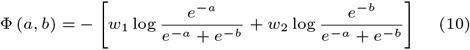

where *w*_1_ and *w*_2_ are both real values in [0,1] and reflect the extent of preference of human experts for the molecules they choose or do not choose. The choice of these two values depend on different tasks and experts’ prior knowledge. Note that the sum of *w*_1_ and *w*_2_ is supposed to be 1.

#### Binary cross entropy loss

Similarly, (10) can be adapted to support the case when only one sample is annotated or samples of identical class are available. In this case, Φ can be a binary cross entropy loss function as follows,

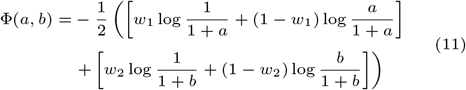

#### Hinge loss

Since our model maps random samples to 3D molecules using straight lines, another intuition to incorporate expert preferences is to train a model that it is easier to find straight lines that are able to generate the samples preferred by experts from random samples, while harder to find a straight line that generates samples not preferred by experts. In order to achieve this, we developed a hinge-like loss function as follows,

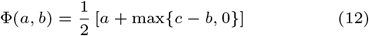

Here *c* is a constant determined by the empirical upper bound of logits of the preferred molecules. Note that Eq. (12) also works for the case when either *a* or *b* is available, which implies that it is not required to provide promising samples and unpromising samples simultaneously. Even if one sample is annotated, our model still can learn the expert knowledge from this sample.

#### Guidance classifier

Recently, Dhariwal & Nichol [7] proposed the classifier guidance technique to boost the sample quality of a diffusion model using an extra trained classifier as guidance. In this work, we also trained a binary classifier with the samples annotated by experts to predict which sample the human experts might select. The training objective of our classifier is defined by

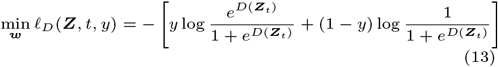

where ***Z***_*t*_ = *t****Z***_1_ + (1 − *t*)***Z***_0_ and *D*(***Z***_*t*_) denotes the logits calculated by the classifier which is parameterized by ***w***. Label *y* is either 0 or 1 with 1 indicating the preferred molecules. We used the gradients of log *p*_***w***_(*y* = 1 | ***Z***_*t*_) with respect to ***Z***_*t*_ to guide the molecule generation process. Specifically, when annotated samples are available, the classifier *D* are jointly trained with all the other neural networks. We denote the refined velocity field by 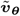. Then we can combine 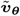_;_ with the classifier *D* to generate molecules as follows,

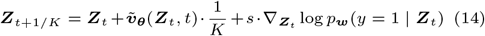

where ***Z***_0_ ∼ 𝒩 (**0, *I***). Note that the number of samples in one preference pair can be greater than 2 for all Φ. The full procedures of human-in-the-loop training and sampling are presented in the Supplementary, section 9.

### Datasets

We trained our ERFM on the training set of the CrossDocked dataset and evaluated its performance related to the basic molecule properties via generating 100 molecules for each protein pocket in the test set (Figure 3 in the main text). In the preference-learning phase, we picked a protein pocket from the test set of the CrossDocked dataset and generate 26,000 molecules for this pocket. According to the preference of interest, among these molecules, we annotated at least 96 molecules as promising drug candidates and at least 96 molecules as unpromising drug candidates (Figure 4 in the main text). The dataset can be found in the key resources table. The details about how we annotated samples can be found in the supplementary materials. Also, the details about data preprocessing can be found in the supplementary materials.

### Simulated preference

In order to evaluate different types of preference, we adopted computer algorithms to play the role of human experts to provide preferences. Specifically, humans can define a threshold for a particular property of interest. If a small molecule generated by AI satisfies the threshold of that property, then this molecule is annotated as a promising sample, or positive sample. Otherwise, it will be annotated as an unpromising one, or negative sample. For instance, one can set the threshold of Vina docking score to −9 to select molecules. In practice, humans may prefer multiple molecular properties. For instance, chemists may prefer small molecules containing benzene rings with reasonable bond lengths. Molecules with benzene rings and reasonable bond lengths are annotated as good samples, while molecules that do not include benzene rings or have unreasonable lengths of bonds are labeled as bad samples.

### Baselines

Both AR [23] and Pocket2Mol [29] are auto-regressive generative models. TargetDiff [11] is a diffusion model which achieves target-aware molecule design by using an SE(3)-equivariant network. It learns continuous atom coordinates and categorical atom types. The results of three baselines are obtained via running their official implementations with default hyper-parameters (see the key resources table for details).

## Supporting information

link4suppofHILDD

## Code availability

The code for reproducing the results reported in the paper is available in the key resources table. For more details regarding implementation, please refer to the supplementary materials.

## Acknowledgments

The authors thank the anonymous reviewers for their valuable suggestions.

---

**Table 1.** Key Resource Table.

| REAGENT or RESOURCE | SOURCE | IDENTIFIER |
| --- | --- | --- |
| **Deposited Data** |  |  |
| CrossDocked Dataset | https://bits.csb.pitt.edu/files/crossdock2020/ | N/A |
| Data Preprocessing | Peng et al.[29] | https://github.com/pengxingang/Pocket2Mol/tree/main/data |
| **Software and Algorithms** |  |  |
| PyTorch | PyTorch | https://pytorch.org/ |
| AR | Luo et al.[23] | https://github.com/luost26/3D-Generative-SBDD |
| Pocket2Mol | Peng et al.[29] | https://github.com/pengxingang/Pocket2Mol |
| TargetDiff | Guan et al.[11] | https://github.com/guanjq/targetdiff |
| Toy experiment | This paper | https://github.com/youmingzhao91/HIL-DD/tree/main/toy-experiment |
| ERFM | This paper | https://github.com/youmingzhao91/HIL-DD |
| HIL-DD | This paper | https://github.com/youmingzhao91/HIL-DD |

## Notes

### Competing Interest Statement

The authors have declared no competing interest.

## References

1. Yuntao Bai, Andy Jones, Kamal Ndousse, Amanda Askell, Anna Chen, Nova DasSarma, Dawn Drain, Stanislav Fort, Deep Ganguli, Tom Henighan, Nicholas Joseph, Saurav Kadavath, Jackson Kernion, Tom Conerly, Sheer El-Showk, Nelson Elhage, Zac Hatfield-Dodds, Danny Hernandez, Tristan Hume, Scott Johnston, Shauna Kravec, Liane Lovitt, Neel Nanda, Catherine Olsson, Dario Amodei, Tom Brown, Jack Clark, Sam McCandlish, Chris Olah, Ben Mann, and Jared Kaplan. Training a helpful and harmless assistant with reinforcement learning from human feedback, 2022.

2. Michael Bostock, Vadim Ogievetsky, and Jeffrey Heer. D3 data-driven documents. IEEE transactions on visualization and computer graphics, 17(12):2301–2309, 2011.

3. Tom B. Brown, Benjamin Mann, Nick Ryder, Melanie Subbiah, Jared Kaplan, Prafulla Dhariwal, Arvind Neelakantan, Pranav Shyam, Girish Sastry, Amanda Askell, et al. Language models are unsupervised multitask learners. OpenAI Blog, 2020.

4. James Chen, Wei Wang, Fan Li, Lei Xiao, Xiaojuan Shen, Lin Fang, Brian L. Bush, and David R. Koes. Dock 6: Impact of new features and current docking performance. Journal of Computational Chemistry, 36(15):1456–1474, 2015.

5. Vijil Chenthamarakshan, Fan Yang, Justin Hoffman, Hendrik Strobelt, Kyunghyun Lim, Martin Wattenberg, Joseph Dunbar, and Erik G Learned-Miller. Cogmol: Target-specific goal-oriented molecular generation. arXiv preprint 2110.08694, 2021.

6. Aakanksha Chowdhery, Sharan Narang, Jacob Devlin, Maarten Bosma, Gabriel Mishra, Adam Roberts, Paul Barham, Hyung Won Chung, Charles Sutton, Simone Gehrmann, et al. Palm: Scaling language modeling with pathways. arXiv preprint 2204.02311, 2022.

7. Prafulla Dhariwal and Alexander Nichol. Diffusion models beat gans on image synthesis. In M. Ranzato, A. Beygelzimer, Y. Dauphin, P.S. Liang, and J. Wortman Vaughan, editors, Advances in Neural Information Processing Systems, volume 34, pages 8780–8794. Curran Associates, Inc., 2021.

8. Paul G Francoeur, Tomohide Masuda, Jocelyn Sunseri, Andrew Jia, Richard B Iovanisci, Ian Snyder, and David R Koes. Three-dimensional convolutional neural networks and a cross-docked data set for structure-based drug design. Journal of chemical information and modeling, 60(9):4200–4215, 2020.

9. Richard A. Friesner, Jay L. Banks, Robert B. Murphy, Thomas A. Halgren, Jasna J. Klicic, D. Todd Mainz, Michael P. Repasky, Edmund H. Knoll, Matthew Shelley, and Jason K. Perry. Extra precision glide: Docking and scoring incorporating a model of hydrophobic enclosure for protein-ligand complexes. Journal of Medicinal Chemistry, 49(21):6177–6196, 2006.

10. Daniel Golovin, John Karro, Greg Kochanski, Chansoo Lee, Xingyou Song, and Qiuyi Zhang. Gradientless descent: High-dimensional zeroth-order optimization. arXiv preprint 1911.06317, 2019.

11. Jiaqi Guan, Wesley Wei Qian, Xingang Peng, Yufeng Su, Jian Peng, and Jianzhu Ma. 3d equivariant diffusion for target-aware molecule generation and affinity prediction. In International Conference on Learning Representations, 2023.

12. Nafisa M Hassan, Amr A Alhossary, Yuguang Mu, and Chee-Keong Kwoh. Protein-ligand blind docking using quickvina-w with inter-process spatio-temporal integration. Scientific reports, 7(1):15451, 2017.

13. Emiel Hoogeboom, Victor Garcia Satorras, Clément Vignac, and Max Welling. Equivariant diffusion for molecule generation in 3d. In International Conference on Machine Learning, pages 8867–8887. PMLR, 2022.

14. Mahmoud Karimi, Haofan Dai, Brian Hie, Fethi Damla Catak, Matteo Neumann, Aniket Arany, Ryan Bressler, and David Becerra. Smiles-bert: Large scale unsupervised pretraining for molecular property prediction. Molecular Pharmaceutics, 19(1):308–318, 2022.

15. Susumu Kobayashi, Titus J Boggon, Tajhal Dayaram, Pasi A Jänne, Olivier Kocher, Matthew Meyerson, Bruce E Johnson, Michael J Eck, Daniel G Tenen, and Balàzs Halmos. Egfr mutation and resistance of non–small-cell lung cancer to gefitinib. New England Journal of Medicine, 352(8):786–792, 2005.

16. Jonas Köhler, Leon Klein, and Frank Noé. Equivariant flows: exact likelihood generative learning for symmetric densities. In International conference on machine learning, pages 5361–5370. PMLR, 2020.

17. Hao Liu, Carmelo Sferrazza, and Pieter Abbeel. Chain of hindsight aligns language models with feedback. arXiv preprint 2302.02676, 2023.

18. Qiang Liu. Rectified flow: A marginal preserving approach to optimal transport. arXiv preprint 2209.14577, 2022.

19. Xingchao Liu, Chengyue Gong, and Qiang Liu. Flow straight and fast: Learning to generate and transfer data with rectified flow. arXiv preprint 2209.03003, 2022.

20. Ilya Loshchilov and Frank Hutter. Cma-es for hyperparameter optimization of deep neural networks, 2016.

21. Ilya Loshchilov and Frank Hutter. Decoupled weight decay regularization. arXiv preprint 1711.05101, 2017.

22. Xiaoyun Lu, Jeff B Smaill, and Ke Ding. Medicinal chemistry strategies for the development of kinase inhibitors targeting point mutations. Journal of Medicinal Chemistry, 63(19):10726–10741, 2020.

23. Shitong Luo, Jiaqi Guan, Jianzhu Ma, and Jian Peng. A 3d generative model for structure-based drug design. In Thirty-Fifth Conference on Neural Information Processing Systems, 2021.

24. Andrea Madotto, Chien-Sheng Wu, and Pascale Fung. Llama: Large language models for question answering. arXiv preprint 2004.07771, 2020.

25. Elman Mansimov, Omar Mahmood, Seokho Kang, and Kyunghyun Cho. Molecular geometry prediction using a deep generative graph neural network. Scientific Reports, 9, 2019.

26. Long Ouyang, Jeffrey Wu, Xu Jiang, Diogo Almeida, Carroll Wainwright, Pamela Mishkin, Chong Zhang, Sandhini Agarwal, Katarina Slama, Alex Ray, John Schulman, Jacob Hilton, Fraser Kelton, Luke Miller, Maddie Simens, Amanda Askell, Peter Welinder, Paul F Christiano, Jan Leike, and Ryan Lowe. Training language models to follow instructions with human feedback. In S. Koyejo, S. Mohamed, A. Agarwal, D. Belgrave, K. Cho, and A. Oh, editors, Advances in Neural Information Processing Systems, volume 35, pages 27730–27744. Curran Associates, Inc., 2022.

27. Junping Pei, Guan Wang, Aoxue Wang, Chengyong Wu, Xiaoli Pan, Wen Shuai, Faqian Bu, Yumeng Zhu, Yuxi Wang, Liang Ouyang, and Weimin Li. Design, synthesis, and antitumor activity of potent and selective egfr l858r/t790m inhibitors and identification of a combination therapy to overcome acquired resistance in models of nonsmall-cell lung cancer. Journal of Medicinal Chemistry, 66(8):5719–5752, 2023. PMID: 37042119.

28. Xingang Peng, Jiaqi Guan, Qiang Liu, and Jianzhu Ma. MolDiff: Addressing the atom-bond inconsistency problem in 3D molecule diffusion generation. In Andreas Krause, Emma Brunskill, Kyunghyun Cho, Barbara Engelhardt, Sivan Sabato, and Jonathan Scarlett, editors, Proceedings of the 40th International Conference on Machine Learning, volume 202 of Proceedings of Machine Learning Research, pages 27611–27629. PMLR, 23–29 Jul 2023.

29. Xingang Peng, Shitong Luo, Jiaqi Guan, Qi Xie, Jian Peng, and Jianzhu Ma. Pocket2mol: Efficient molecular sampling based on 3d protein pockets. In International Conference on Machine Learning, 2022.

30. Alec Radford, Karthik Narasimhan, Tim Salimans, and Ilya Sutskever. Improving language understanding by generative > pre-training. OpenAI blog, 1(2), 2018.

31. Dhwani Raghav, Vinay Sharma, and Subhash Mohan Agarwal. Structural investigation of deleterious non-synonymous snps of egfr gene. Interdisciplinary Sciences: Computational Life Sciences, 5:60–68, 2013.

32. Victor Garcia Satorras, Emiel Hoogeboom, and Max Welling. E (n) equivariant graph neural networks. In International conference on machine learning, pages 9323–9332. PMLR, 2021.

33. Arne Schneuing, Yuanqi Du, Charles Harris, Arian Jamasb, Ilia Igashov, Weitao Du, Tom Blundell, Pietro Lió, Carla Gomes, Max Welling, et al. Structure-based drug design with equivariant diffusion models. arXiv preprint 2210.13695, 2022.

34. LLC Schrödinger. Schrödinger Suite, 2021. Version 7.0.

35. David Sehnal, Sebastian Bittrich, Mandar Deshpande, Radka Svobodová, Karel Berka, Václav Bazgier, Sameer Velankar, Stephen K Burley, Jaroslav Koča, and Alexander S Rose. Mol* viewer: modern web app for 3d visualization and analysis of large biomolecular structures. Nucleic acids research, 49(W1):W431–W437, 2021.

36. Jiayi Shen, Tao Zhang, Su-Jie Zhu, Min Sun, Linjiang Tong, Mengzhen Lai, Rong Zhang, Wei Xu, Ruibo Wu, Jian Ding, Cai-Hong Yun, Hua Xie, Xiaoyun Lu, and Ke Ding. Structure-based design of 5-methylpyrimidopyridone derivatives as new wild-type sparing inhibitors of the epidermal growth factor receptor triple mutant (egfrl858r/t790m/c797s). Journal of > Medicinal Chemistry, 62(15):7302–7308, 2019. PMID: 31298540.

37. Richard A Stein and James V Staros. Insights into the evolution of the erbb receptor family and their ligands from sequence analysis. BMC evolutionary biology, 6(1):1–17, 2006.

38. Zhiwei Tang, Dmitry Rybin, and Tsung-Hui Chang. Zeroth-order optimization meets human feedback: Provable learning via ranking oracles, 2023.

39. Oliver Trott and Arthur J. Olson. Autodock vina: Improving the speed and accuracy of docking with a new scoring function, efficient optimization, and multithreading. Journal of Computational Chemistry, 31(2):455–461, 2010.

40. Marcel L. Verdonk, Jason C. Cole, Michael J. Hartshorn, Christopher W. Murray, and Richard D. Taylor. Gold 5.2: Ligand docking with selectivity, specificity, and exposure constraints. Journal of Medicinal Chemistry, 47(24):6188– 6196, 2004.

41. Robert C. Wade, Rafik R. Gabdoulline, and Win D. Wade. Moe: A computer program for molecular operating environment. Journal of Chemical Information and Modeling, 30(2):377–385, 1990.

42. Minkai Xu, Lantao Yu, Yang Song, Chence Shi, Stefano Ermon, and Jian Tang. Geodiff: A geometric diffusion model for molecular conformation generation. In International Conference on Learning Representations, 2022.

43. Wei Zhou, Xiaofeng Liu, Zhengchao Tu, Lianwen Zhang, Xin Ku, Fang Bai, Zhenjiang Zhao, Yufang Xu, Ke Ding, and Honglin Li. Discovery of pteridin-7(8h)-one-based irreversible inhibitors targeting the epidermal growth factor receptor (egfr) kinase t790m/l858r mutant. Journal of Medicinal Chemistry, 56(20):7821–7837, 2013. PMID: 24053674.

