## Supplementary material for "The Human-In-the-Loop Drug Design Framework with Equivariant Rectified Flow": link4suppofHILDD

[1]Department of Mathematics, Technical University of Darmstadt, Karolinenpl. 5, 64289, 64289, Hesse, Germany, [2]State Key Laboratory of Natural and Biomimetic Drugs, School of Pharmaceutical Sciences, Peking University, 38 Xueyuan Road, 100191, Beijing, China, [3]Institute for Artificial Intelligence, Peking University, 5 Yiheyuan Road, 100871, Beijing, China, [4]School of Intelligence Science and Technology, Peking University, 5 Yiheyuan Road, 100871, Beijing, China, [5]PersLab Research, PersLab, Tianjin, China, [6]Department of Computer Science, University of Texas at Austin, 2317 Speedway, 78712, Texas, USA, [7]Department of Computer Science, Indiana University at Bloomington, Luddy Hall, 700 N Woodlawn Ave, 47408, Indiana, USA, [8]Helixon US Inc, USA, [9]Department of Electronic Engineering, Tsinghua University, 30 Shuangqing Road, 100084, Beijing, China and [10]Institute for AI Industry Research, Tsinghua University, Zhongguancun East Road No.1-8 Block C, 100084, Beijing, China

#### Abstract

This article contains all the technical details for reproducing the work *The Human-In-the-Loop Drug Design Framework with Equivariant Rectified Flow.*

#### A. A note on translation equivariance

Schneuing et al. [19] demonstrates that the subspace trick is not required for 3D model constructions. The main reason is that in order to achieve translation equivariance, we can translate a Gaussian noise to the center of the pocket prior to training the model. This translation is equivalent to moving the center of the pocket to the origin for both training and sampling. At the end of the generation process, the generated molecule is moved back to the center of the pocket. It is important to note that while $\boldsymbol{x}_0$ needs to be centered at the origin, it is not necessary to perform this operation on $\boldsymbol{x}_t$, as the neural network will learn the relative distance from $\boldsymbol{x}_t$ during the training process.

#### B. Toy experiment for preference learning

Given a rectified flow (RF) model represented by the velocity field $\boldsymbol{v_\theta}$, obtained by fitting $\boldsymbol{X} = (\boldsymbol{X}_0, \boldsymbol{X}_1)$, where $\boldsymbol{X}_0$ represents random Gaussian noise and $\boldsymbol{X}_1$ represents the observed data, we utilized $\boldsymbol{v_\theta}$ to generate samples $\boldsymbol{Z}_1$ by applying Gaussian noise $\boldsymbol{Z}_0$. Let $S^+$ denote samples that satisfy specific properties, and $S^-$ denote samples that do not satisfy these properties. These samples can be divided into two types: good samples $\boldsymbol{Z}_1^+ \in S^+$ with corresponding noise $\boldsymbol{Z}_0^+$, and bad samples $\boldsymbol{Z}_1^- \in S^-$ with corresponding noise $\boldsymbol{Z}_0^-$. Let $\boldsymbol{Z}^+ = (\boldsymbol{Z}_1^+, \boldsymbol{Z}_0^+)$ and $\boldsymbol{Z}^- = (\boldsymbol{Z}_1^-, \boldsymbol{Z}_0^-)$, then a good sample and a bad sample form a pair of proposals $(\boldsymbol{Z}^+, \boldsymbol{Z}^-) \in P$, where $P$ represents the set of proposals. Using these proposals, we optimize the following objective function over proposals $(\boldsymbol{Z}^+, \boldsymbol{Z}^-) \in \boldsymbol{P}$.

$$\min_{\boldsymbol{\theta}} \ell_{pref}(\boldsymbol{X}, \boldsymbol{Z}^+, \boldsymbol{Z}^-, t) = \ell(\boldsymbol{X}, t) + \lambda \cdot \Phi\left(\ell(\boldsymbol{Z}^+, t), \ell(\boldsymbol{Z}^-, t)\right) \tag{1}$$

where $t \in [0, 1]$ and $\lambda$ is a positive scalar. The loss function (1) consists of multiple components, each with its own effect on the molecule generation process as follows:

1. $\ell(\boldsymbol{X}, t)$: It represents the training loss of the RF model on the observed data $\boldsymbol{X}$. Its purpose is to encourage the model to learn the distribution of the observed data.
2. $\ell(\boldsymbol{Z}^+, t)$ and $\ell(\boldsymbol{Z}^-, t)$: These terms represent the straightness of the velocity field $\boldsymbol{v_\theta}$ at $t$ on the generated samples, $\boldsymbol{Z}^+$ (positive samples) and $\boldsymbol{Z}^-$ (negative samples), respectively. Minimizing $\ell(\boldsymbol{Z}^+, t)$ can make the velocity field generate the positive samples more easily, which encourages the model to capture the specific patterns and characteristics associated with the positive samples. On the other hand, maximizing $\ell(\boldsymbol{Z}^-, t)$ makes the velocity field generate the negative in a harder way, which helps the model to avoid the undesirable patterns represented by the negative samples. By incorporating these loss functions, the model can generate samples that align better with the desired distribution.
3. $\Phi(a, b)$: This is a function designed to decrease $a$ and increase $b$ during the optimization process, in which case $a = \ell(\boldsymbol{Z}^+, t)$ and $b = \ell(\boldsymbol{Z}^-, t)$. Three types of $\Phi(a, b)$ have been proposed in the main text. It helps in efficiently learning desirable features by scoring the samples differently based on their desirability.
4. $\lambda$: This is a hyper-parameter determining the weight of the penalty term, which indicates the importance of human preferences in the fine-tuning process. We need to choose an appropriate $\lambda$ to ensure the penalty term does not dominate the fine-tuning process. We employed an automatic and harmless regularization algorithm proposed by [8] and $\alpha = \beta = 1$ in their algorithm.

By jointly optimizing (1), a pretrained model is fine-tuned to generate samples that not only resemble the desired distribution but also exhibit characteristics that differentiate them from the undesirable molecules. The fine-tuning process aligns the generated samples with human preferences, avoids undesirable features, and leads to the production of better samples that more closely approximate the desired distribution.

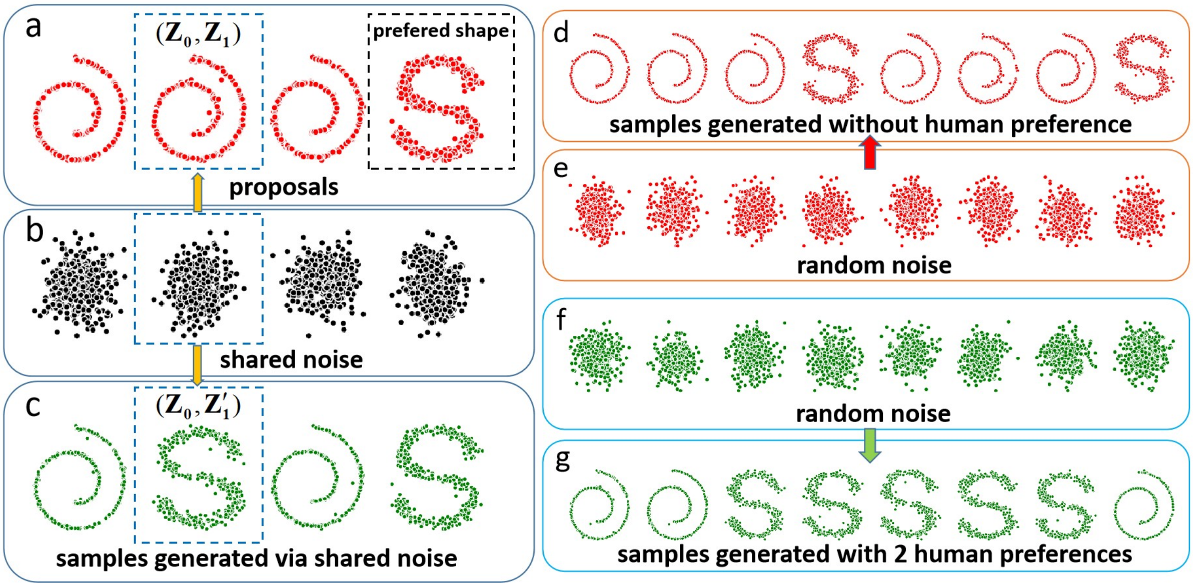


**Fig. 1.** A toy experiment demonstrating the preference learning.

We applied our methodology on a toy dataset to validate the hypothesis described above. The dataset consisted of two types of samples: Swiss roll and S curve, which were generated using the functions make_s_curve and make_swiss_roll from the sklearn.datasets library [16]. We first trained a RF model, denoted as *model A*, to generate samples. The generated samples were then annotated by a human expert based on their preference for either selecting the Swiss rolls or S curves. We then fine-tuned a copy of *model A* using the selected samples, which yields another model, named *model B*. It's worth noting that *model A* tends to generate more Swiss roll samples without any human interference.

Initially, *Model A* generates only four samples as depicted in Figure 1. Out of these samples, only the S curve sample is preferred. Using these four samples along with the provided annotation, *Model B* learns this preference by optimizing equation (1). Consequently, *Model B* is updated and transforms the same set of noise shown in Figure 1b into four samples shown in Figure 1c. Notably, the second noise pattern, which was originally converted into a Swiss roll by *Model A*, is successfully transformed into an S curve by *Model B*. This serves as evidence for the effectiveness of optimizing equation (1). Remarkably, this type of improvement is typically achievable within just three updates, highlighting the efficiency of our algorithm.

To evaluate the learning performance of our algorithm, we provided both *model A* and *model B* with two sets of random noise, as shown in Figures 1e and 1f. Consequently, *model B* generated a significantly larger number of S curves (Figure 1f) compared to the samples generated by *model A* (Figure 1d). With only a few preferences and updates, *model B* successfully corrects the bias of *model A*. The code used to construct the toy dataset and these models are available at: `https://github.com/youmingzhao91/HIL-DD/tree/main/toy-experiment`.

#### C. Results for other four pockets

In this section, we present the performance of the proposed HIL-DD for proteins 14GS, 4YHJ, 5W2G, and 1COY. As shown in Figures 2-5, The results are similar to the results presented in Figure 4 in the main article.

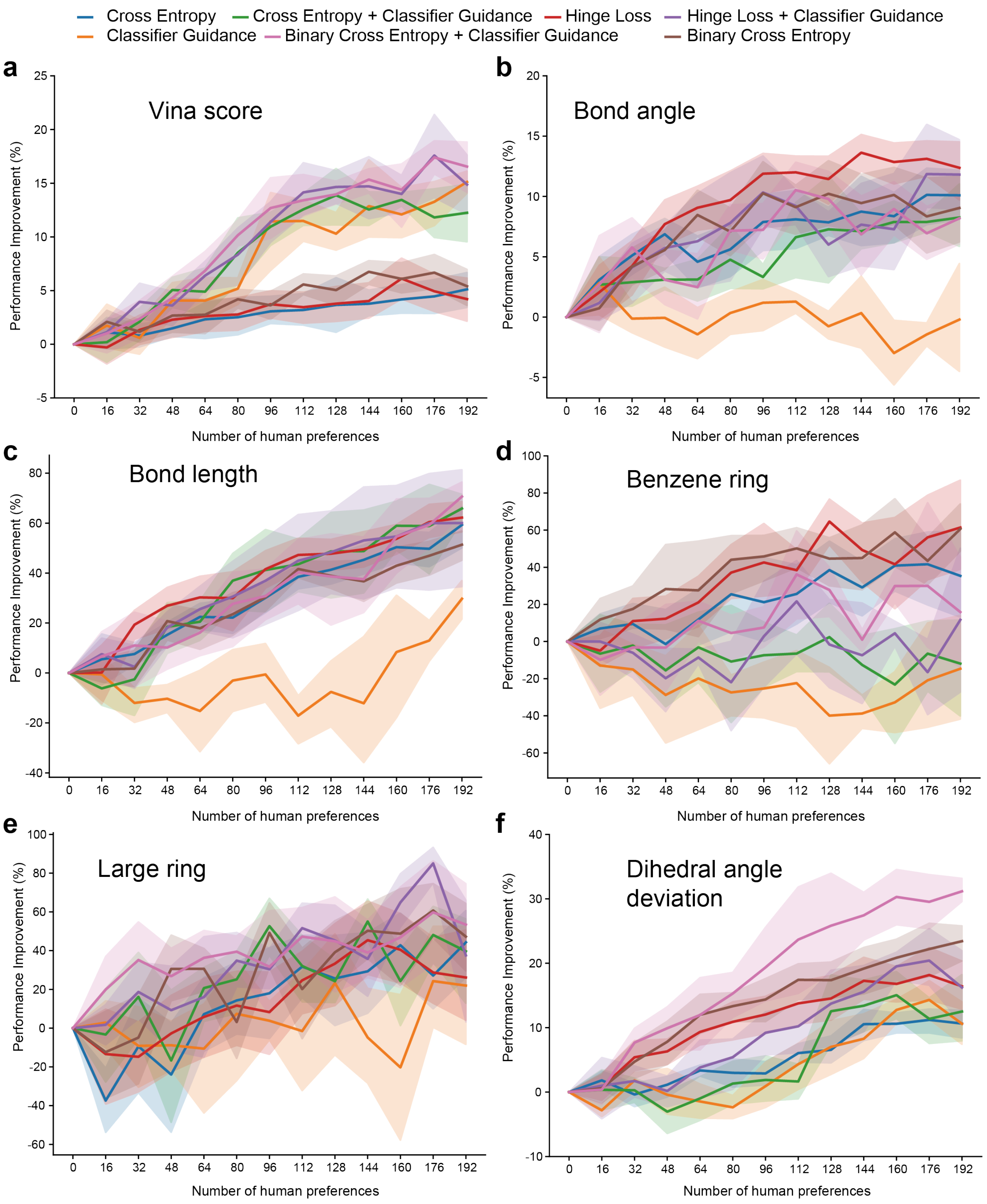


**Fig. 2.** Performance of protein 14GS

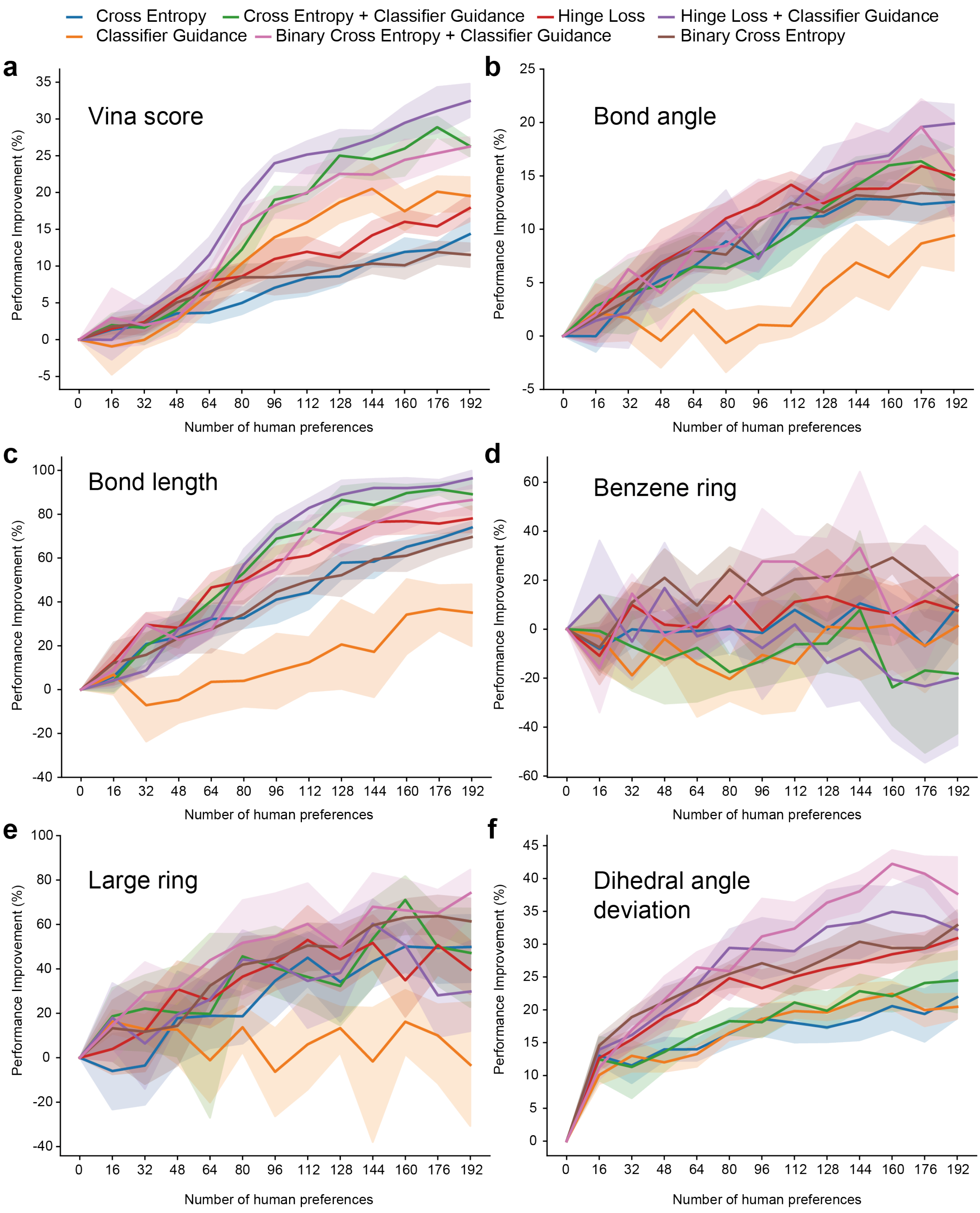


**Fig. 3.** Performance of protein 4YHJ

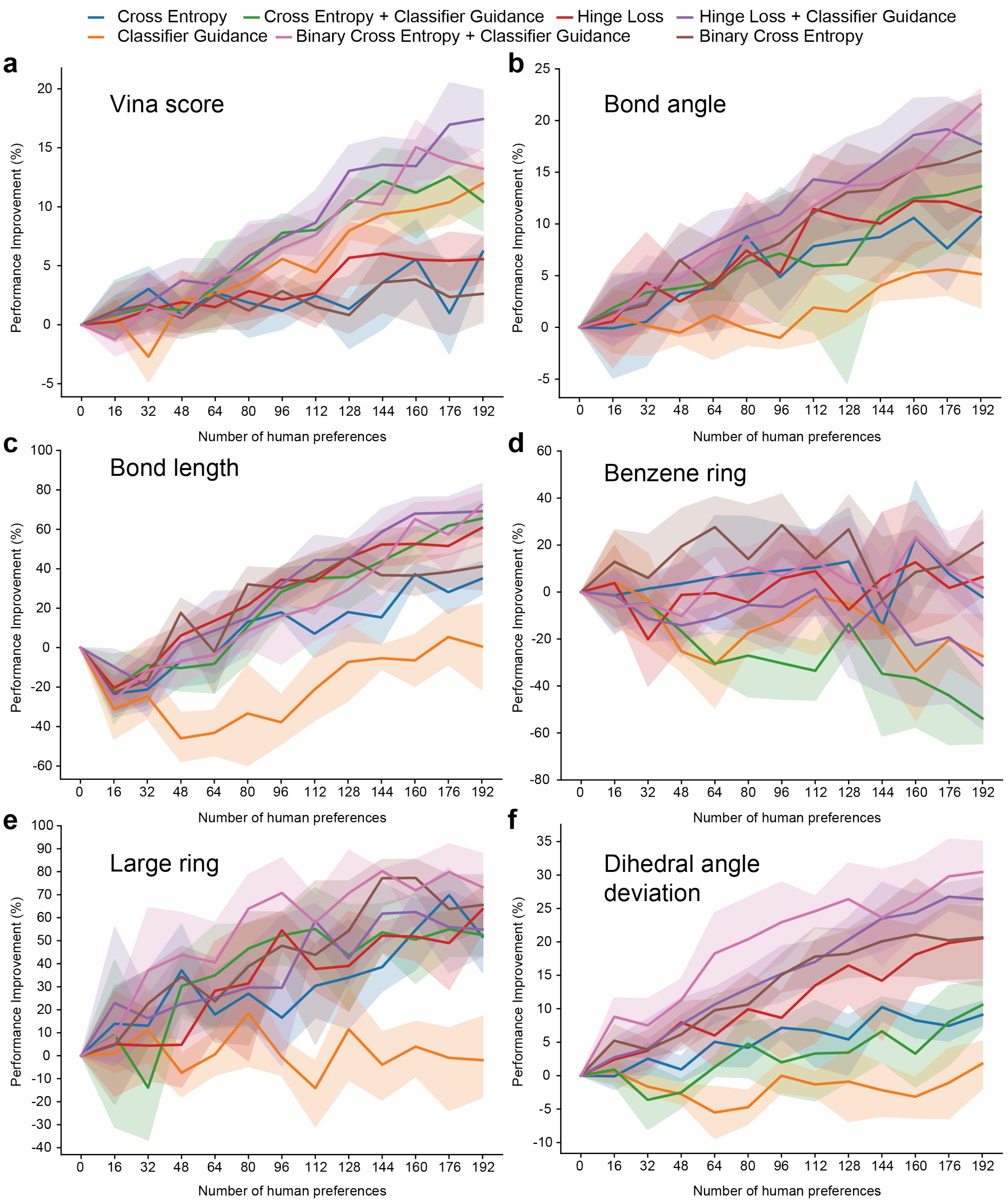


**Fig. 4.** Performance of protein 5W2G

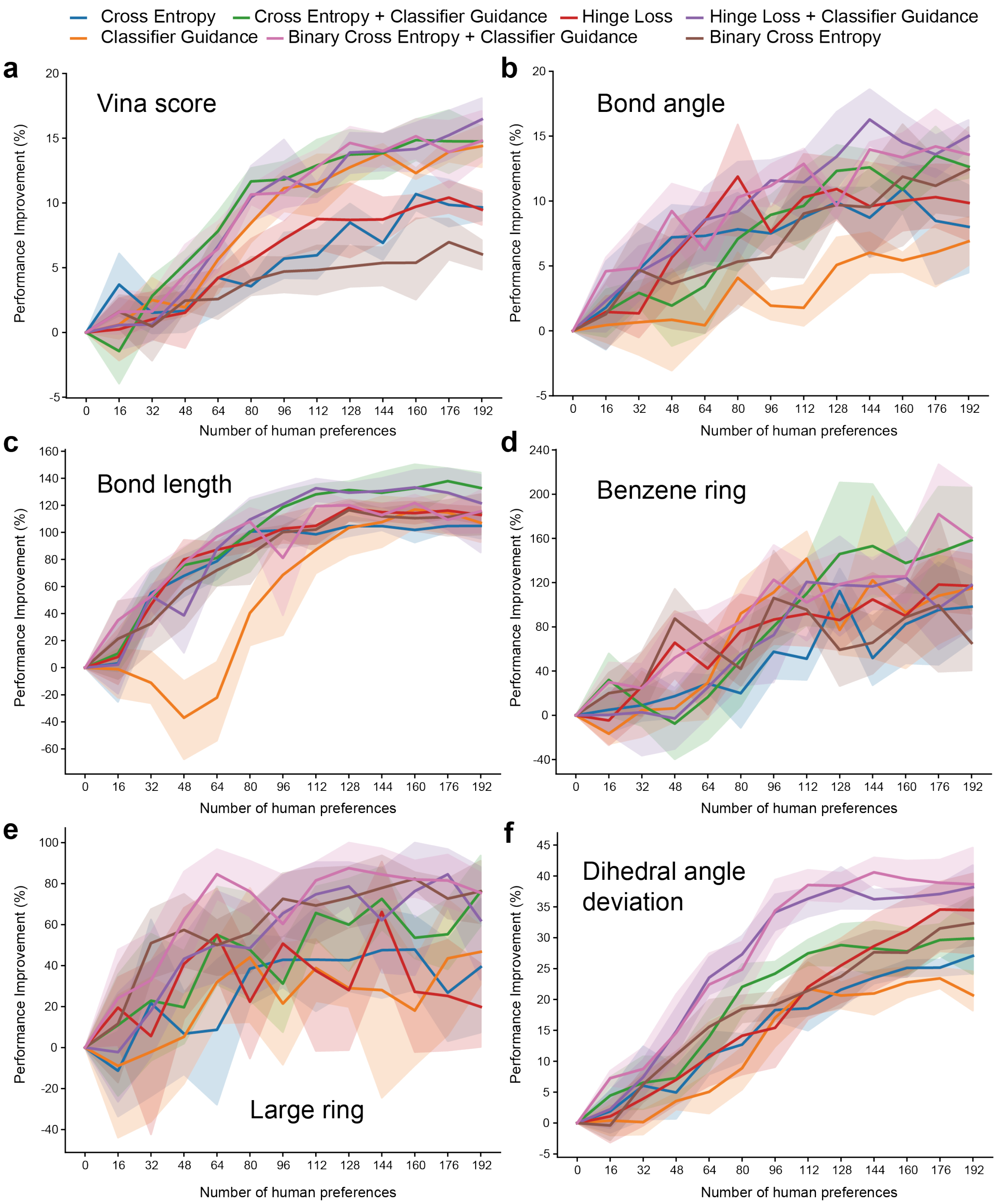


Fig. 5. Performance of protein 1COY

### D. Model construction

#### D.1. Feature embedding

The features of a 3D protein pocket consist of the atom type embedding $\boldsymbol{v}_h^P \in \mathbb{R}^{16}$, the amino acid type embedding $\boldsymbol{v}_{aa}^P \in \mathbb{R}^{16}$, and an embedding $\boldsymbol{v}_{ib}^P \in \mathbb{R}^8$ indicating whether an atom belongs to backbone of an amino acid. The features of a small molecule comprise the atom type embedding $\boldsymbol{v}_h \in \mathbb{R}^{32}$, the bond type embedding $\boldsymbol{v}_b \in \mathbb{R}^{32}$, and the time embedding $t \in \mathbb{R}^8$. We used a linear layer to project the combination of all features and time embeddings to new feature vectors that have the common hidden dimension of the EGNN module.

$$\boldsymbol{h}^P = \text{Linear}(\text{concat}[\boldsymbol{v}_h^P, \boldsymbol{v}_{aa}^P, \boldsymbol{v}_{ib}^P]) \tag{2}$$

$$\boldsymbol{h} = \text{Linear}(\text{concat}[\boldsymbol{v}_h, t]) \tag{3}$$

$$\boldsymbol{b} = \text{Linear}(\text{concat}[\boldsymbol{v}_b, t]) \tag{4}$$

#### D.2. Edge encoder

In the **Method** section, the edge function $\phi_e$ is defined as follows:

$$\boldsymbol{m}_{ij} = \text{MLP}(\boldsymbol{h}_i^l, \boldsymbol{h}_j^l, d_{ij}^2, a_{ij}) \tag{5}$$

where $d_{ij}$ represents the Euclidean distance between atoms $i$ and $j$:

$$d_{ij} = \|\boldsymbol{x}_i^l - \boldsymbol{x}_j^l\|_2^2 \tag{6}$$

In addition, an extra attribute $a_{ij} = \|\boldsymbol{x}_i^0 - \boldsymbol{x}_j^0\|_2^2$ is included at the first layer. Note that $a_{ij}$ is preserved for all EGNN modules. The last layer of the MLP employs the SiLU activation [4] instead of a linear layer.

The attention function $\phi_{\text{att}}$ takes the message $\boldsymbol{m}_{ij}$ as input and produces a scalar $\tilde{e}_{ij} \in (0, 1)$ as follows:

$$\tilde{e}_{ij} = \text{Sigmoid}(\text{Linear}(\boldsymbol{m}_{ij})) \tag{7}$$

#### D.3. Node encoder

Node embedding function $\phi_h$ takes the features of atom $i$ and aggregates its neighbor information and outputs new representations of atom $i$ as follows,

$$\text{MLP}(\boldsymbol{h}_i, \sum_{j \in \mathcal{N}(i)} \tilde{e}_{ij}\boldsymbol{m}_{ij}) \tag{8}$$

where $\boldsymbol{h}_i$ indicates the features of the $i$-th atom of the small molecule. Note that the neighbors $\mathcal{N}(i)$ of a node $i$ on the small molecule can be an atom from a protein pocket.

#### D.4. Equivariant model

The equivariant model $\phi_x$ has the same inputs as $\phi_e$ but outputs a scalar.

$$\text{MLP}(\boldsymbol{h}_i, \boldsymbol{h}_j, d_{ij}, a_{ij}) \tag{9}$$

where MLP ends up with a linear layer whose output dimension is 1. Finally, we apply a linear layer to the predicted velocity field for all molecule coordinates. This linear layer only have one weight. Note that there is no weight for bias of the linear layer to preserve equivariance [3].

$$\boldsymbol{x} = w\boldsymbol{x}. \tag{10}$$

#### D.5. Bond model

The bond model $\phi_b$ takes the same inputs as $\phi_e$ but $a_{ij}$ should be replaced by $\boldsymbol{b}_{ij}$ and outputs the updated bond features.

$$\text{MLP}(\boldsymbol{h}_i, \boldsymbol{h}_j, d_{ij}^2, b_{ij}^l). \tag{11}$$

**Table 1.** Evaluation of generated molecules for 100 protein pockets of the CrossDocked test set. The numbers following ERFM represent the number of time steps when sampling molecules.

| | Vina Score (↓) | QED (↑) | SA (↑) | Lipinski (↑) | Diversity (↑) | Time (s, ↓) |
|---|---|---|---|---|---|---|
| Test set | $-6.87 \pm 2.30$ | $0.48 \pm 0.20$ | $0.73 \pm 0.14$ | $4.34 \pm 1.14$ | – | – |
| ERFM-1000 | $-6.87 \pm 1.64$ | $0.43 \pm 0.14$ | $0.62 \pm 0.09$ | $4.55 \pm 0.61$ | $0.78 \pm 0.06$ | $334.99 \pm 103.48$ |
| ERFM-100 | $-6.88 \pm 1.65$ | $0.43 \pm 0.14$ | $0.62 \pm 0.09$ | $4.56 \pm 0.62$ | $0.78 \pm 0.06$ | $34.36 \pm 10.85$ |
| ERFM-50 | $-6.90 \pm 1.88$ | $0.43 \pm 0.20$ | $0.62 \pm 0.12$ | $4.55 \pm 0.81$ | $0.78 \pm 0.06$ | $17.32 \pm 5.50$ |
| ERFM-20 | $-6.89 \pm 1.86$ | $0.43 \pm 0.20$ | $0.62 \pm 0.12$ | $4.56 \pm 0.79$ | $0.77 \pm 0.06$ | $7.16 \pm 2.36$ |
| ERFM-10 | $-6.92 \pm 1.86$ | $0.42 \pm 0.20$ | $0.61 \pm 0.12$ | $4.55 \pm 0.76$ | $0.77 \pm 0.05$ | $3.69 \pm 1.23$ |
| ERFM(w/o-bond) | $-6.64 \pm 1.75$ | $0.37 \pm 0.20$ | $0.60 \pm 0.11$ | $4.38 \pm 0.96$ | $0.79 \pm 0.06$ | $333.73 \pm 104.12$ |
| HIL-DD | $-7.64 \pm 2.00$ | $0.46 \pm 0.19$ | $0.63 \pm 0.13$ | $4.75 \pm 0.57$ | $0.78 \pm 0.06$ | $332.80 \pm 125.22$ |

##### D.6. The effect of sampling with different numbers of time steps

Since our ERFM learned an ODE to follow straight paths connecting Gaussian noise and molecules of the training data, which implies that the learned ODE can be possibly simulated accurately with coarse time discretization in the sampling phase [12]. To validate this, the researchers utilized the ERFM to sample molecules using different numbers of uniformly spaced time steps: 1000, 100, 50, 20, and 10. The results, shown in Table 1, indicate that reducing the number of time steps does not compromise the quality of the generated molecules while enabling faster sampling. For instance, sampling 100 molecules with only 10 time steps took an average of 3.69 seconds and achieved better Vina scores.

##### D.7. The effect of training ERFM with/without incorporating bond information

To examine the impact of incorporating bond information during training on model performance, we conducted an ablation study by training the ERFM model with and without bond information and comparing the results.

As shown in Table 1, incorporating bond information during training improves performance across all evaluation metrics except for diversity. This indicates that providing the model with explicit bond information is beneficial for optimizing binding affinity (Vina score), drug-likeness (QED and Lipinski score), and synthetic accessibility (SA score). Meanwhile, their performance on diversity is comparable. Overall, this ablation study demonstrates that incorporating bond information is important for generating molecules with improved pharmacological properties, without sacrificing diversity.

##### D.8. The effect of fine-tuning for sampling performance

To assess the generalization of the HIL-DD model, the researchers conducted multiple preference evaluations simultaneously only on PDB 2V3R. These preferences included Vina score, QED, and SA. The preference thresholds for Vina score were set at -9 and -8 for good and bad molecules, respectively. For QED, the thresholds were 0.65 for good molecules and 0.5 for bad molecules, and for SA, the thresholds were 0.65 for good molecules and 0.6 for bad molecules. To identify positive samples, molecules that satisfied all three criteria (Vina score, QED, and SA) were selected. Conversely, negative samples were chosen from molecules that failed to satisfy any of the three criteria. The labels for positive samples were determined using a linear combination of the Vina score, QED, and SA, i.e. $-0.1 \times$ Vina score + QED + SA. The labels for negative samples are 0.

The ERFM was fine-tuned through 5 runs, following the procedure outlined in Section E.3. In each run, out of 12 evaluations, the model with the best performance in terms of Vina score, QED, and SA was selected as the final HIL-DD model. This final model, chosen from the 5 runs, was then used to sample 100 molecules for each protein pocket in the CrossDocked test set. As shown in Table 1, the HIL-DD model outperforms ERFM-1000 significantly in terms of Vina score, while also demonstrating improvements in QED and SA, though the HIL-DD model had never seen the other 99 protein pockets in the test set. The demonstrates that HIL-DD has great generalization ability.

#### E. Hyper-parameters of HIL-DD

##### E.1. Hyper-parameters of the neural networks

Our Equivariant Rectified Flow Model (ERFM) is composed of five layers of equivariant graph convolutional nerual networks (EGNNs) and feature-mapping layers before and after the EGNN module. The features of the original data are mapped to 40-d vectors, which has been described in D.1. The number of hidden features in EGNN is 256. After combining 32-d bond embeddings with 8-d time embeddings, we projected the 40-d bond features to 256-d feature vectors before the EGNN module. Following the EGNN module, all features are mapped to 32-d features as the outputs, except for atomic coordinates which remain 3-d throughout ERFM. The number of trainable parameters is about 4.4 million. Note that all parameters for constructing embeddings remain fixed in the training stage.

##### E.2. Hyper-parameters of training

We employed the optimizer AdamW to train our model and the batch size is 1. In the first 20,000 updates, we warmed up the learning rate from 0 to 1e-3 in a linear way. The parameters are smoothed by exponential moving average (EMA) as conducted in [20], with a factor of 0.9999. We evaluated our model on the test set and decayed the learning rate by a factor of 0.8 if the validation loss did not decrease five times. The training process takes about two days on Nvidia GTX 4090 GPU.

##### E.3. Hyper-parameters of sampling

In the molecule generation step, the molecule size is the same as the size of the reference molecule in the dataset. Our model supports bond generation by learning the velocity field for bond types. However, we still use Open Babel [14] to predict the bond types when reconstructing a 3D molecule since Open Babel gives more reasonable bond types.

##### E.4. Hyper-parameters of fine-tuning

We conducted fine-tuning of the trained ERFM model using the AdamW optimizer with a learning rate of 0.0004 and a weight decay of 0.001. Our fine-tuning process considered 6 preferences: Vina score, bond angle, bond length, benzene ring, large ring, and dihedral angle deviation. To ensure comprehensive evaluation, we performed 5 rounds of our algorithm for each preference.

Within each round, we provided the model with 8 positive annotations and 8 negative annotations a total of 12 times, which we refer to as "12 injections of preferences." All annotations were stored in memory throughout each run, and we reused all previous annotations for the remainder of the round. After each injection, we evaluated the current model by generating samples and calculating the metrics of interest. For Vina score, the number of generated samples was 20, while for all other metrics, it was 100.

The number of model updates was evenly spaced within the range of 50 to 200 across the 12 injections for each round. It should be noted that the range for the preference of avoiding large rings was [100,1000]. In each update, we fine-tuned the model using 2 positive samples and 2 negative samples. It is important to mention that new annotations in memory were sampled with a slightly higher probability since old annotations had been used multiple times.

When incorporating classifier guidance into the fine-tuning process, the guiding strength was consistently set at 0.1.

#### F. Classifier guidance model

##### F.1. Model architecture

The classifier guidance model adopts a structure comprising two graph convolutional layers (GCLs) along with additional feature mapping layers before and after the GCLs. The entire process is outlined by Eqs (12) to (16). The input components consist of atomic coordinates, element-type embeddings, and bond-type embeddings. Notably, the use of coordinates solely pertains to distance computations, which nicely empowers the classifier with rotational invariance and translational invariance properties. The output consists of 1-dimensional logits, serving as the input for the binary cross-entropy loss function. The classifier model is defined by,

$$\boldsymbol{h} = \text{Linear}(\text{concat}[\boldsymbol{h}, t]) \tag{12}$$

$$\boldsymbol{e}_{ij} = \text{concat}([d_{ij}, \boldsymbol{b}_{ij}, t]) \tag{13}$$

$$\boldsymbol{m}_{ij} = \phi_e(\boldsymbol{h}_i^{(l)}, \boldsymbol{h}_j^{(l)}, \boldsymbol{e}_{ij}) \tag{14}$$

$$\boldsymbol{h}_i^{(l+1)} = \phi_h\left(\boldsymbol{h}_i^{(l)}, \sum_{j \in \mathcal{N}(i)} \boldsymbol{m}_{ij}\phi_{\text{att}}(\boldsymbol{m}_{ij})\right) \tag{15}$$

Here $\phi_e$ is defined by Eq (5), where $d_{ij}$ represents the distance between atoms encoded by radial basis function (RBF) kernels [17]. $\phi_{att}$ and $\phi_h$ are defined by Eqs (7) and (8), respectively. Notably, the classification model considers the eight nearest neighbors when constructing the graph. Following this, we apply a linear layer to map $\boldsymbol{h}$ to 32-dimensional vectors, and subsequently perform a global-mean operation on $\boldsymbol{h} \in \mathbb{R}^{n\times 32}$ across $n$ atoms. This yields the final representation of $\boldsymbol{h} \in \mathbb{R}^{32}$. The scalar output (logits) is obtained using a linear layer in accordance with Eq (16) as follows,

$$\text{logits} = \text{Linear}(\text{global\_mean}(\text{Linear}(\boldsymbol{h}))) \tag{16}$$

##### F.2. Determining the scale of guiding strength

In order to determine an appropriate scale for the guiding strength, we conducted tests using different values: 0.0001, 0.001, 0.01, 0.05, 0.1, 0.2, and 1. We found that as the scale of the guiding strength increased, the performance of the relevant metric of interest also improved. However, if the guiding strength was too strong, it led to a decrease in atom stability.

Based on our observations, a guiding strength scale of 0.1 provided a good balance between performance improvement and atom stability. This particular value allowed for significant enhancement in the relevant metric of interest while avoiding excessive negative effects on atom stability.

#### G. Technical details for preference learning

We sampled 26,000 molecules conditioned on each pocket taken from the CrossDocked test set, including 2V3R, 14GS, 4YHJ, 5W2G, and 1COY. We conducted comprehensive experiments for 6 preferences, including Vina score, bond angle, bond length, benzene ring, large ring, and dihedral angle deviation.

**Table 2.** Statistics of 6 preferences for PDB 2V3R.

| | Mean | Threshold of unpromising ligands | Threshold of promising ligands | Best improved mean | Maximum improvement(%) |
|---|---|---|---|---|---|
| Vina score | −7.46 | −8.00 | −10.00 | −9.88 ± 0.18 | 32.43 ± 2.47 |
| Bond angle | −6.67 | −4.00 | −3.50 | −5.40 ± 0.17 | 18.99 ± 2.50 |
| Bond length | −4.64 | 0.00 | 1.00 | 0.72 ± 0.13 | 115.51 ± 2.88 |
| Benzene ring | 2.78% | - | c1ccccc1 | 43.69% ± 3.44% | 1471.67 ± 123.78 |
| Large ring | 7.20% | 9-membered ring or larger ring | 6-membered ring or smaller ring | 0.17% ± 0.34% | 96.30 ± 7.60 |
| Dihedral angle deviation | 17.13 | 10.00 | 5.00 | 8.68 ± 1.04 | 59.34 ± 4.86 |

**Table 3.** Statistics of 6 preferences for PDB 14GS.

| | Mean | Threshold of unpromising ligands | Threshold of promising ligands | Best improved mean | Maximum improvement(%) |
|---|---|---|---|---|---|
| Vina score | −6.78 | −7.00 | −9.00 | −7.97 ± 0.29 | 17.59 ± 4.34 |
| Bond angle | −6.67 | −5.00 | −4.00 | −6.10 ± 0.18 | 13.63 ± 2.62 |
| Bond length | −4.64 | 0.00 | 1.50 | −1.42 ± 0.34 | 70.63 ± 6.97 |
| Benzene ring | 15.68% | - | c1ccccc1 | 25.81% ± 3.14% | 64.63 ± 20.00 |
| Large ring | 4.48% | 9-membered ring or larger ring | 6-membered ring or smaller ring | 1.30% ± 0.91% | 85.01 ± 10.47 |
| Dihedral angle deviation | 21.36 | 10.00 | 5.00 | 11.73 ± 0.34 | 31.16 ± 2.00 |

**Table 4.** Statistics of 6 preferences for PDB 4YHJ.

| | Mean | Threshold of unpromising ligands | Threshold of promising ligands | Best improved mean | Maximum improvement(%) |
|---|---|---|---|---|---|
| Vina score | −7.64 | −8.00 | −10.00 | −9.20 ± 0.38 | 20.30 ± 4.97 |
| Bond angle | −7.22 | −5.00 | −4.50 | −5.78 ± 0.17 | 19.91 ± 2.33 |
| Bond length | −4.34 | 0.00 | 1.00 | −0.59 ± 0.33 | 96.32 ± 4.77 |
| Benzene ring | 21.83% | - | c1ccccc1 | 29.09% ± 8.19% | 33.23 ± 37.50 |
| Large ring | 13.07% | 9-membered ring or larger ring | 6-membered ring or smaller ring | 3.38% ± 1.61% | 74.12 ± 12.30 |
| Dihedral angle deviation | 17.39 | 10.00 | 5.00 | 10.03 ± 0.43 | 42.24 ± 2.49 |

##### G.1. Vina score

For the selected molecules, the labels are determined based on the preference loss. The labels for the unselected molecules are set to 0. In the case of cross entropy loss, the labels for the unselected molecules are obtained by applying a softmax function to their Vina scores. This allows a more nuanced representation of the preference for unselected molecules, considering their Vina scores as a measure of binding affinity. For hinge loss and binary cross entropy loss, the labels of the selected samples are normalized relative to the best binding affinity within a pair. The labels of the selected molecules with the best binding affinity in a pair are set to 1, while the labels of other selected molecules are assigned values in the range of [0.6,1). Here, we introduce a lower bound of 0.6 for positive molecules to ensure a certain level of confidence in their selection. This allows for a range of label values to capture the varying degrees of human preference within the set of selected molecules.

**Table 5.** Statistics of 6 preferences for PDB 5W2G.

| | Mean | Threshold of unpromising ligands | Threshold of promising ligands | Best improved mean | Maximum improvement(%) |
|---|---|---|---|---|---|
| Vina score | $-6.96$ | $-7.00$ | $-8$ | $-8.18 \pm 0.19$ | $17.43 \pm 2.79$ |
| Bond angle | $-7.41$ | $-5.00$ | $-4.50$ | $-5.81 \pm 0.11$ | $21.56 \pm 1.50$ |
| Bond length | $-4.64$ | 0.00 | 0.50 | $-1.28 \pm 0.37$ | $72.46 \pm 7.91$ |
| Benzene ring | 23.73% | - | c1ccccc1 | $30.48\% \pm 3.56\%$ | $28.46 \pm 14.99$ |
| Large ring | 14.87% | 9-membered ring or larger ring | 6-membered ring or smaller ring | $2.93\% \pm 0.11\%$ | $80.30 \pm 0.77$ |
| Dihedral angle deviation | 14.77 | 10.00 | 5.00 | $10.27 \pm 0.78$ | $30.45 \pm 5.29$ |

**Table 6.** Statistics of 6 preferences for PDB 1COY.

| | Mean | Threshold of unpromising ligands | Threshold of promising ligands | Best improved mean | Maximum improvement(%) |
|---|---|---|---|---|---|
| Vina score | $-8.21$ | $-8.00$ | $-10.50$ | $-9.56 \pm 0.15$ | $16.45 \pm 1.88$ |
| Bond angle | $-6.09$ | $-5.00$ | $-3.50$ | $-5.10 \pm 0.15$ | $16.28 \pm 2.50$ |
| Bond length | $-1.73$ | 0.00 | 1.50 | $-1.55 \pm 0.5$ | $137.90 \pm 11.43$ |
| Benzene ring | 12.17% | - | c1ccccc1 | $28.08\% \pm 7.7\%$ | $181.94 \pm 49.14$ |
| Large ring | 6.27% | 9-membered ring or larger ring | 6-membered ring or smaller ring | $0.78\% \pm 0.96\%$ | $87.54 \pm 15.26$ |
| Dihedral angle deviation | 20.37 | 10.00 | 5.00 | $12.10 \pm 0.50$ | $40.59 \pm 2.45$ |

##### G.2. Bond angle

In this work, we focused on ten most common bond angle types found in the CrossDocked training set: ccc, CCC, Ccc, CCO, CCN, CNC, cnc, Occ, CC=O, and C=CC. In this context, lowercase letters represent aromatic bonds. We used a Gaussian Mixture Model (GMM) to fit the distribution of specific bond angle values within the CrossDocked dataset. For each bond angle type, we utilized the GaussianMixture function from the sklearn.mixture library [16] to learn and fit the distribution.

##### G.3. Bond length

For the preference learning of bond lengths, our experiment focuses on the ten most common bond types: cc, CC, CN, CO, cn, C=O, OP, C=C, CF, and O=P. For each bond type, Gaussian Mixture Model (GMM) is employed to fit the distribution of its bond lengths within the CrossDocked dataset. The GaussianMixture module from the sklearn.mixture library [16] is utilized for learning the distribution associated with each bond type. The molecules demonstrating high average log likelihood are selected as positive samples, while the molecules with low average log likelihood are considered as negative samples.

##### G.4. Benzene ring

The selection of molecules based on benzene rings depends on two key factors. Firstly, molecules that exhibit a high average log likelihood of favorable bond lengths are preferred. Specifically, in our implementation, we set the threshold of 1.0 for the average log likelihood of bond lengths as criterion. This emphasizes the importance of favorable bond lengths in the formation of a benzene ring, as it requires six identical aromatic bonds. Secondly, only molecules that contain benzene rings are considered. Thus, only those molecules that satisfy both criteria are selected as promising ones.

##### G.5. Large ring

Typically, human experts tend to avoid molecules that include large rings, such as those with nine or more atoms. Therefore, we selected molecules containing only small rings composed of less than seven atoms as positive samples. Conversely, molecules consisting solely of large rings composed of more than eight atoms are considered negative samples.

##### G.6. Dihedral angle deviation

Selecting a metric to measure the preference of dihedral angles can be challenging due to the presence of irregular spikes in certain dihedral angle types in the training set. It becomes difficult for the Gaussian mixture model (GMM) to capture these outliers effectively. Instead of using log likelihood, we focused on identifying typical dihedral angle modes for each dihedral angle type. After excluding those with highly irregular dihedral angle modes, we arrived at the ten most common dihedral angle types listed in Table 7. We then computed the deviations between the real dihedral angles and the typical values for all generated molecules. Finally, we selected samples with low average dihedral angle deviations as positive samples, while samples with higher deviations were classified as negative samples.

**Table 7.** 10 common dihedral angle types.

| dihedral angle types | modes(°) |
|---|---|
| cc-cc-cc | -180, 0, 180 |
| CC-CC-CC | -180, -60, 0, 60, 180 |
| cc-cc-CC | -180, 0, 180 |
| cc-cn-nc | -180, 0, 180 |
| cc-cc-cn | -180, 0, 180 |
| cn-nc-cn | -180, 0, 180 |
| OC-cc-cc | -180, 0, 180 |
| cc-cc-CN | -180, 0, 180 |
| O=C-CN-NC | -180, 0, 180 |
| CC-CC-C=C | -180, -135, 0, 135, 180 |

#### H. Training and molecule generation algorithms of ERFM

In this section, the detailed algorithmic procedures for training and molecule generation of ERFM are outlined in Algorithm 1 and Algorithm 2, respectively.

**Algorithm 1** ERFM training algorithm

1: **Input:** Protein-molecule complex data $\mathcal{D}$, neural network $\boldsymbol{v_\theta}$ and learning rate $\alpha$.
2: **repeat**
3: Sample $(\boldsymbol{Z}_1, \boldsymbol{P}) \sim \mathcal{D}$, $\boldsymbol{Z}_0 \sim \mathcal{N}(\mathbf{0}, \boldsymbol{I})$, and $t \in \mathcal{U}(0,1)$
4: Subtract center of gravity from $\boldsymbol{Z}_0$
5: Compute $\boldsymbol{Z}_t = (1-t)\boldsymbol{Z}_0 + t\boldsymbol{Z}_1$
6: Minimize $\ell(\boldsymbol{Z}, t)$ via

$$\boldsymbol{\theta} \leftarrow \boldsymbol{\theta} - \alpha \cdot \nabla_{\boldsymbol{\theta}} \ell(\boldsymbol{Z}, t)$$

7: **until** converged
8: **Output:** Trained velocity flow model $\boldsymbol{v_\theta}$.

**Algorithm 2** ERFM molecule generation algorithm

1: **Input:** Protein pocket $\boldsymbol{P}$, pre-trained velocity field $\boldsymbol{v_\theta}$, and number of training steps $K$.
2: Sample $\boldsymbol{Z}_0 \sim \mathcal{N}(\mathbf{0}, \boldsymbol{I})$
3: Subtract center of gravity from $\boldsymbol{Z}_0$
4: **for** $k = 0, 1, \cdots, K-1$ **do**
5: $\boldsymbol{Z}_{(k+1)/K} = \boldsymbol{Z}_{k/K} + \boldsymbol{v_\theta}\left(\boldsymbol{Z}_{k/K}, k/K\right) \cdot \frac{1}{K}$
6: **end for**
7: $\boldsymbol{Z}_1 = \boldsymbol{Z}_{(K-1+1)/K}$
8: **return** $\boldsymbol{Z}_1$

#### I. Training and sampling algorithms of HIL-DD

In this section, the detailed algorithms for training the HIL-DD framework by using the preference learning are described in Algorithm 3 and Algorithm 4, respectively.

**Algorithm 3** HIL-DD training algorithm

**Input:** Protein-molecule complex data $\mathcal{D}$, protein pocket $\boldsymbol{P}$, and pre-trained velocity field $\boldsymbol{v}_{\boldsymbol{\theta}}$.
**Input:** No. of positive samples in one pair $k^+$ and No. of negative samples in one pair $k^-$, and No. of updates $m$.
Generate ligand set $\mathcal{S} = \{\boldsymbol{Z} = (\boldsymbol{Z}_0, \boldsymbol{Z}_1)\}$ conditioned on $\boldsymbol{P}$ via Algorithm 2.
Experts make a promising ligand set $\mathcal{S}^+$ and an unpromising ligand set $\mathcal{S}^-$.
**for** $i = 1, 2, \cdots, m$ **do**
    Sample $\boldsymbol{X} = (\boldsymbol{X}_0, \boldsymbol{X}_1)$, where $\boldsymbol{X}_0 \sim \mathcal{N}(\boldsymbol{0}, \boldsymbol{I})$ and $(\boldsymbol{X}_1, \boldsymbol{P}) \sim \mathcal{D}$
    Pick $k^+$ promising ligands $\mathcal{Z}^+ = \{\boldsymbol{Z}^+ \mid \boldsymbol{Z}^+ \in \mathcal{S}^+\}$
    Pick $k^-$ unpromising ligands $\mathcal{Z}^- = \{\boldsymbol{Z}^- \mid \boldsymbol{Z}^- \in \mathcal{S}^-\}$
    Sample $t \in \mathcal{U}(0, 1)$
    Minimize the classification loss via

$$\boldsymbol{w} \leftarrow \boldsymbol{w} - \alpha \cdot \nabla_{\boldsymbol{w}} \left( \frac{1}{k^+} \sum_{\boldsymbol{Z}^+ \in \mathcal{Z}^+} \ell_D(\boldsymbol{Z}^+, t, 1) + \frac{1}{k^-} \sum_{\boldsymbol{Z}^- \in \mathcal{Z}^-} \ell_D(\boldsymbol{Z}^-, t, 0) \right)$$

    Minimize $\ell_{pref}(\boldsymbol{X}, \boldsymbol{Z}^+, \boldsymbol{Z}^-, t)$ via

$$\boldsymbol{\theta} \leftarrow \boldsymbol{\theta} - \alpha \cdot \nabla_{\boldsymbol{\theta}} \ell_{pref}(\boldsymbol{X}, \boldsymbol{Z}^+, \boldsymbol{Z}^-, t)$$

**end for**
**return** Refined velocity field $\tilde{\boldsymbol{v}}_{\boldsymbol{\theta}}$ and trained classifier $D$.

**Algorithm 4** Human-in-the-loop ligand sampling conditioned on a protein pocket

**Input:** protein pocket $\boldsymbol{P}$, and either refined velocity field $\tilde{\boldsymbol{v}}^{\boldsymbol{\theta}}$ or classifier $D$ with gradient scale $s$, or both of them.
Sample $\boldsymbol{Z}_0 \sim \mathcal{N}(\boldsymbol{0}, \boldsymbol{I})$
**for** $t = 0, 1/K, 2/K, \cdots, (K-1)/K$ **do**
    $\boldsymbol{Z}_{t+1/K} = \boldsymbol{Z}_t + \tilde{\boldsymbol{v}}^{\boldsymbol{\theta}}(\boldsymbol{Z}_t, t) \cdot \frac{1}{K} + s \cdot \nabla_{\boldsymbol{Z}_t} \log p_{\boldsymbol{w}}(y = 1 \mid \boldsymbol{Z}_t)$
**end for**
**return** $\boldsymbol{Z}_1$

#### J. More molecule visualization

In Figure 6, more molecules are visualized for the experiments corresponding to Fig. 4c,e,f of the main text. Note that the molecules residing in the same column of each panel are generated with the same noise. As shown in Figure 6, HIL-DD can convert a noise that was originally transported to a bad molecule into a good molecule, which implies that our HIL-DD has the ability to expand the chemical space of promising drug candidates.

#### K. Data preprocessing

We adopted the CrossDocked dataset [7] as the training and test data. The dataset initially contains 22.5 million poses of small molecules docked into proteins, among which there are 2,922 unique pockets and 13,839 unique small molecules. We adopted the filtered and processed non-redundant subsets of the CrossDocked dataset as previous works [13, 18, 9], which contains 100,000 and 100 protein-molecule pairs for training and test, respectively. After removing hydrogens for all small molecules, we further processed the data as follows. A protein pocket is represented by its atom coordinates, atom types, amino acid types associated with the atoms, and a binary variable to indicate whether an atom belongs to the backbone or side-chain of an amino acid. All features are represented as embeddings except atom coordinates. The dimensions of the embeddings are 16, 16, and 8, respectively. A small molecule is represented by its atom coordinates and atom types. There are seven atom types that appear in the pre-processed data. We considered the aromatic information of all atoms of the small molecule, which involves the Carbon, Nnitrogen, Oxygen, Phosphorus, and Sulphur. The bond types we considered include: single, double, triple, aromatic bonds, and unspecified. Atomic types and chemical bond types information are represented as two separate 32-dimensional embeddings. These embeddings, along with an additional 8-dimensional embedding representing time, are concatenated together to form the final input of our ERFM.

#### L. Implementation details

Our HIL-DD is implemented in Python 3.10 and PyTorch 1.12 [15], along with other packages including Numpy 1.23.3 [10], SciPy 1.9.1 [21], PyTorch Geometric 2.1.0 [6], and PyTorch Scatter 2.0.9 [5]. We used some open-source tools OpenBabel 3.1.1 [14, 2] and RDKit [1] to calculate relevant metrics, e.g., QED, SA, Lipinski, and diversity. Qvina2 [11] is employed to calculate Vina docking scores. We utilized the AdamW optimizer with a learning rate of 1e-3 and a weight decay of 1e-3 to train the ERFM. The batch size is set to 1. We decayed the learning rate by a factor of 0.8 when the validation loss plateaus. It takes about 48 hours to train

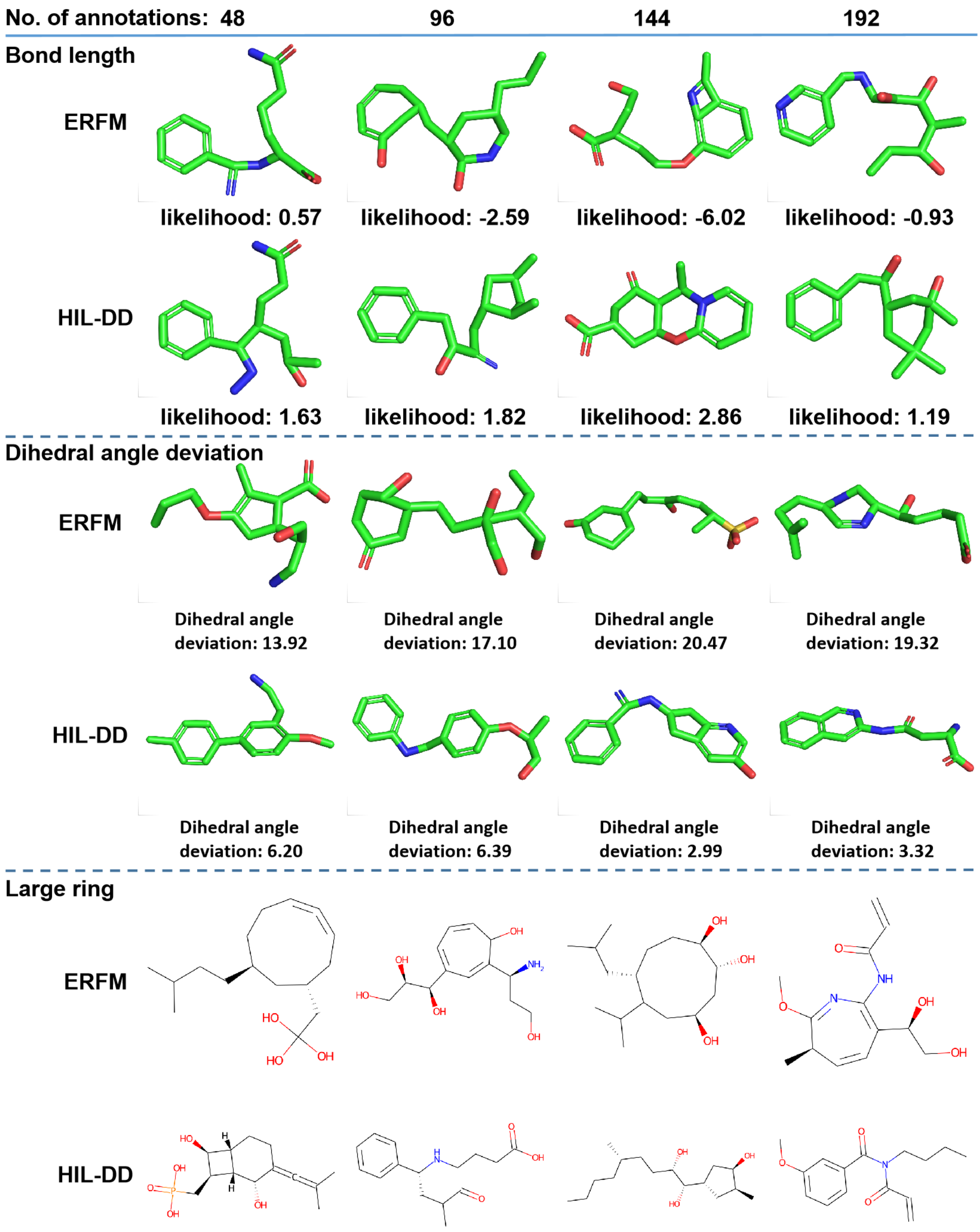


**Fig. 6.** Drug candidates generated by ERFM and HIL-DD when conducting the experiments corresponding to Fig. 4c,e,f of the main text. The molecules are shown in three panels. In each panel, the molecules in each column are generated with the same noise. In the second row of each panel, the molecules presented in 4 columns are generated by finetuning the pretrained ERFM with 48, 96, 144, and 192 annotations, respectively. The likelihood is measured in a log scale.

the ERFM on one NVIDIA GeForce RTX 4090 GPU card. We employed the AdamW optimizer with a learning rate of 4e-5 and a weight decay of 1e-3 to refine the ERFM with preference learning.

#### Acknowledgments

The authors thank the anonymous reviewers for their valuable suggestions.

#### References

1. RDKit: Open-source cheminformatics. `https://www.rdkit.org`, 2023.
2. The Open Babel Package, version 3.1.1. `https://openbabel.org/wiki/Main_Page`, 2023.
3. Congyue Deng, Or Litany, Yueqi Duan, Adrien Poulenard, Andrea Tagliasacchi, and Leonidas J. Guibas. Vector neurons: A general framework for so(3)-equivariant networks. In *Proceedings of the IEEE/CVF International Conference on Computer Vision (ICCV)*, pages 12200–12209, October 2021.
4. Stefan Elfwing, Eiji Uchibe, and Kenji Doya. Sigmoid-weighted linear units for neural network function approximation in reinforcement learning. *Neural networks*, 107:3–11, 2018.
5. Matthias Fey. PyTorch Extension Library of Optimized Scatter Operations. `https://pypi.org/project/torch-scatter/`, 2023.
6. Matthias Fey and Jan E. Lenssen. Fast graph representation learning with PyTorch Geometric. In *ICLR Workshop on Representation Learning on Graphs and Manifolds*, 2019.
7. Paul G Francoeur, Tomohide Masuda, Jocelyn Sunseri, Andrew Jia, Richard B Iovanisci, Ian Snyder, and David R Koes. Three-dimensional convolutional neural networks and a cross-docked data set for structure-based drug design. *Journal of chemical information and modeling*, 60(9):4200–4215, 2020.
8. Chengyue Gong, Xingchao Liu, and Qiang Liu. Automatic and harmless regularization with constrained and lexicographic optimization: A dynamic barrier approach. In M. Ranzato, A. Beygelzimer, Y. Dauphin, P.S. Liang, and J. Wortman Vaughan, editors, *Advances in Neural Information Processing Systems*, volume 34, pages 29630–29642, 2021.
9. Jiaqi Guan, Wesley Wei Qian, Xingang Peng, Yufeng Su, Jian Peng, and Jianzhu Ma. 3d equivariant diffusion for target-aware molecule generation and affinity prediction. In *International Conference on Learning Representations*, 2023.
10. Charles R. Harris, K. Jarrod Millman, Stéfan J. van der Walt, Ralf Gommers, Pauli Virtanen, David Cournapeau, Eric Wieser, Julian Taylor, Sebastian Berg, Nathaniel J. Smith, Robert Kern, Matti Picus, Stephan Hoyer, Marten H. van Kerkwijk, Matthew Brett, Allan Haldane, Jaime Fernández del Río, Mark Wiebe, Pearu Peterson, Pierre Gérard-Marchant, Kevin Sheppard, Tyler Reddy, Warren Weckesser, Hameer Abbasi, Christoph Gohlke, and Travis E. Oliphant. Array programming with NumPy. *Nature*, 585(7825):357–362, September 2020.
11. Nafisa M Hassan, Amr A Alhossary, Yuguang Mu, and Chee-Keong Kwoh. Protein-ligand blind docking using quickvina-w with inter-process spatio-temporal integration. *Scientific reports*, 7(1):15451, 2017.
12. Xingchao Liu, Chengyue Gong, and Qiang Liu. Flow straight and fast: Learning to generate and transfer data with rectified flow. *arXiv preprint arXiv:2209.03003*, 2022.
13. Shitong Luo, Jiaqi Guan, Jianzhu Ma, and Jian Peng. A 3d generative model for structure-based drug design. In *Thirty-Fifth Conference on Neural Information Processing Systems*, 2021.
14. Noel M O'Boyle, Michael Banck, Craig A James, Chris Morley, Tim Vandermeersch, and Geoffrey R Hutchison. Open babel: An open chemical toolbox. *Journal of cheminformatics*, 3(1):1–14, 2011.
15. Adam Paszke, Sam Gross, Soumith Chintala, Gregory Chanan, Edward Yang, Zachary DeVito, Zeming Lin, Alban Desmaison, Luca Antiga, and Adam Lerer. Automatic differentiation in pytorch. In *NIPS-W*, 2017.
16. F. Pedregosa, G. Varoquaux, A. Gramfort, V. Michel, B. Thirion, O. Grisel, M. Blondel, P. Prettenhofer, R. Weiss, V. Dubourg, J. Vanderplas, A. Passos, D. Cournapeau, M. Brucher, M. Perrot, and E. Duchesnay. Scikit-learn: Machine learning in Python. *Journal of Machine Learning Research*, 12:2825–2830, 2011.
17. Xingang Peng, Jiaqi Guan, Qiang Liu, and Jianzhu Ma. Moldiff: Addressing the atom-bond inconsistency problem in 3d molecule diffusion generation. *arXiv preprint arXiv:2305.07508*, 2023.
18. Xingang Peng, Shitong Luo, Jiaqi Guan, Qi Xie, Jian Peng, and Jianzhu Ma. Pocket2mol: Efficient molecular sampling based on 3d protein pockets. In *International Conference on Machine Learning*, 2022.
19. Arne Schneuing, Yuanqi Du, Charles Harris, Arian Jamasb, Ilia Igashov, Weitao Du, Tom Blundell, Pietro Lió, Carla Gomes, Max Welling, et al. Structure-based drug design with equivariant diffusion models. *arXiv preprint arXiv:2210.13695*, 2022.
20. Yang Song and Stefano Ermon. Improved techniques for training score-based generative models. *Advances in neural information processing systems*, 33:12438–12448, 2020.
21. Pauli Virtanen, Ralf Gommers, Travis E. Oliphant, Matt Haberland, Tyler Reddy, David Cournapeau, Evgeni Burovski, Pearu Peterson, Warren Weckesser, Jonathan Bright, Stéfan J. van der Walt, Matthew Brett, Joshua Wilson, K. Jarrod Millman, Nikolay Mayorov, Andrew R. J. Nelson, Eric Jones, Robert Kern, Eric Larson, C J Carey, İlhan Polat, Yu Feng, Eric W. Moore, Jake VanderPlas, Denis Laxalde, Josef Perktold, Robert Cimrman, Ian Henriksen, E. A. Quintero, Charles R. Harris, Anne M. Archibald, Antônio H. Ribeiro, Fabian Pedregosa, Paul van Mulbregt, and SciPy 1.0 Contributors. SciPy 1.0: Fundamental Algorithms for Scientific Computing in Python. *Nature Methods*, 17:261–272, 2020.
